# Historical genomics reveals the evolutionary mechanisms behind multiple outbreaks of the host-specific coffee wilt pathogen *Fusarium xylarioides*

**DOI:** 10.1101/2020.08.07.241695

**Authors:** D. Peck, R. W. Nowell, J. Flood, M. J. Ryan, T. G. Barraclough

## Abstract

Nearly 50% of crop yields are lost to pests and disease, with plants and pathogens locked in an amplified co-evolutionary process of disease outbreaks. Coffee wilt disease, caused by *Fusarium xylarioides*, decimated coffee production in west and central Africa following an initial 1920s outbreak. After successful management, it later re-emerged reaching two separate epidemics by the 2000s on arabica coffee in Ethiopia and robusta coffee in east and central Africa. Here, we use genome sequencing of six historical culture collection strains spanning 70 years to identify the evolutionary processes behind these repeated outbreaks. The robusta population arose from the initial outbreak, whilst the arabica population is divergent and emerged independently. The two populations evolved similar pathologies by separately acquiring different effector genes horizontally via transposable elements from other *Fusarium* taxa, including *F. oxysporum*. Thus, historical genomics can help reveal mechanisms that allow fungal pathogens to keep pace with humanity’s efforts to resist them.

## 2 Introduction

Fungal diseases have devastated major crop yields throughout history and continue to do so (Fones et al. 2017, Strange & Scott 2005). Large-scale planting of crops generates strong selection for new pathogens to emerge, which leads to further rounds of plant breeding to develop new resistant genotypes. This leads to “boom and bust cycles” that intensify the natural co-evolutionary dynamics of hosts and pathogens. A key goal for sustainable agriculture is therefore to predict disease outbreaks and design robust evolutionary solutions for long-term protection (Burdon & Thrall 2008). A first step towards this goal is to understand the genetic and evolutionary mechanisms by which pathogens overcome resistance and infect new host species. Plants have innate defence responses to detect and overcome pathogen attack (Jones & Dangl 2006). In response, an emerging pathogen can evolve new mechanisms to suppress and overwhelm basal plant defences. These could arise by mutation (including gene duplication or loss), recombination and selection operating within a single population, or from hybridization and/or horizontal gene transfer between species to generate new pathogenicity variants. Strong selection to evade plant immunity also leads to host-specificity, whereby pathogens evolve to target particular species or varieties (Sanchez-Vallet et al. 2018). For example, *Fusarium oxysporum*’s well-studied host-specific *formae speciales* (f. sp.) cause disease on over 120 plant species, including Panama disease of bananas, *F. oxysporum* f. sp. *cubense* (Michielse & Rep 2009).

Comparative genomics is revealing the mechanisms that promote rapid evolution of effector proteins and host specificity in fungal pathogens. Effector genes are often found in highly mutable parts of the genome (de Jonge et al. 2011). For example, in ascomycete fungi effectors often occupy AT-rich compartments of the genome with high mutation rates or cluster with transposable elements (TEs), which increase variation via duplications, deletions, insertions and inversions (Ma et al. 2010, Klosterman et al. 2011, Rouxel et al. 2011, Chuma et al. 2011, Schmidt et al. 2013). In addition, many ascomycetes have mechanisms to facilitate horizontal transfer of effector genes between taxa either by “pathogenicity islands” in which pathogenicity genes and TEs cluster in chromsosomal segments depleted in GC (Han et al. 2001) or by whole mobile chromosomes carrying suites of effectors. For instance, the ability of *F. oxysporum f. sp.* lycopersici to infect tomatoes derives from a lineage-specific mobile chromosome, which can be transferred experimentally between strains (Ma et al. 2010). Pathogenicity can therefore evolve by mutation, recombination and selection operating within a single lineage, or from horizontal gene transfer between strains to generate new pathogenicity variants.

Although comparative genomics has uncovered mechanisms behind host specialisation in several fungal pathogens, exactly how these processes play out during disease cycles remains less clear. Studies have mostly compared contemporary lineages with different host specialisations, rather than tracking genetic changes over time. For example, understanding the roles of within-lineage evolution versus horizontal transfer in generating new effector gene combinations would benefit from comparing genomes before and after boom-bust cycles, as well as between differentially adapted host-specialists.

Here, we take advantage of historic strains collected over the last 70 years to investigate successive outbreaks and the origin of host specialisation in *Fusarium xylarioides* Steyaert, a soil-borne fungal pathogen that causes coffee wilt disease (CWD). CWD first emerged as a devastating disease of *Coffea excelsa* and *C. canephora* crops in west and central Africa from the 1920s to 1960s (figure 1). Improved crop sanitation and breeding programmes successfully reduced its impact but CWD later re-emerged in the 1970s, spreading extensively throughout the 1990s and 2000s. It now comprises two host-specific and geographically separated populations, one on *C. canephora* robusta coffee in Uganda, Tanzania and DRC and the other on *C. arabica* in Ethiopia (figure 1). Both populations cause significant losses to the coffee cash crop, on Africa’s two most valuable species (Olal et al. 2018, Mulatu & Shanko 2019). *F. xylarioides* therefore offers a unique study system with repeated epidemics and the emergence of two host-specific populations (Lepoint et al. 2005, Flood 2005, 2009). Critically, historical living strains are optimally cryopreserved in a living state in culture collections that come from the earlier pre-1970s outbreaks as well as more recent strains.

**Figure 1:**
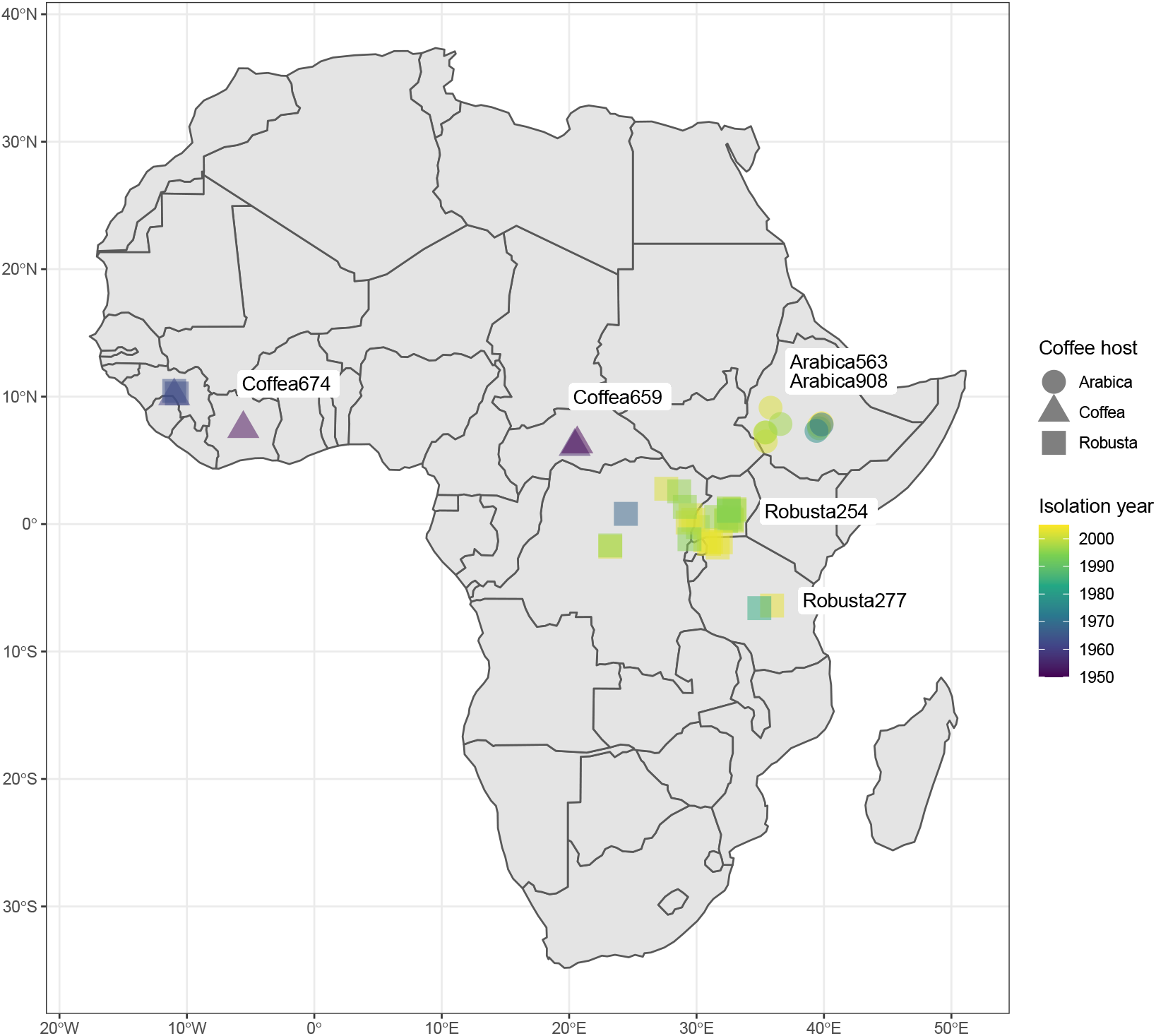
The emergence of *F. xylarioides*. A map of Africa detailing the year collected, country of origin and coffee plant host for 62 *F. xylarioides* strains in the CABI fungal culture collection. Few strains were collected pre-1970s, hence the small number detailed here

Previous work described the pathology of the epidemics (Flood 2009), clarified molecular taxonomy (Buddie et al. 2015, Flood 2005), showed reproductive isolation between the host-specialists (Lepoint et al. 2005), and reported the first genome (Olal et al. 2019). The genetic basis for successive outbreaks and host-specialisation remains unexplored, however. Wilting occurs when a pathogen proliferates in and blocks the host xylem, so restricting water transport (Tjamos & Beckman 1989, Mace et al. 1981). In order to colonize the xylem vascular system, effector proteins, including carbohydrate-active enzymes such as cellulases and pectinases, are required by the fungus to degrade and penetrate the root system (Sharma et al. 2016, Clérivet et al. 2000). In *F. oxysporum*, effector proteins behind wilt induction (termed SIX for Secreted In Xylem) are encoded by a single mobile, pathogenicity chromosome (Ma et al. 2010, Schmidt et al. 2013). As a result, the same host-specific f. sp.’s can have polyphyletic origins, as the ability to infect a particular host is transferred horizontally (Abo et al. 2005, Ellis et al. 2014, 2016, O’Donnell et al. 1998). Whether similar mechanisms apply for *F. xylarioides* and CWD, and how pathogenicity is restored between successive disease outbreaks, remains unknown. Intriguingly, coffee is intercropped with banana, and *F. xylarioides* and *F. oxysporum* have been co-isolated from roots of both plants in Uganda, and from coffee roots in Ethiopia (Flood 2005, Serani et al. 2007). Indeed, *F. oxysporum* is able to infect coffee, where it induces a wilt but does not result in the trees’ death (Serani et al. 2007). These findings raise the possibility that *F. xylarioides* may have acquired certain pathogenicity genes from *F. oxysporum*, that has facilitated the recent outbreaks on coffee.

To address these questions, we sequenced and compared the genomes of six representative historical *F. xylarioides* strains from the CABI-IMI living culture collection: two encompassing the initial pre-1970s outbreak (termed “*Coffea”)*, two from the 1990-2000s robusta population (termed “robusta”), and two from the 1990-2000s arabica population (termed “arabica”). Each pair of strains was originally collected five years apart and from different countries (except the arabica population, which is only found in Ethiopia, figure 1). Current evidence from molecular markers and crossing experiments supports the distinctiveness of the arabica and robusta populations, but varies with respect to relationships between them and with the initial outbreak’s *Coffea* strains. Differing studies show the 1990-2000s arabica and robusta populations as sister clades (Lepoint 2006), or the 1990-2000s robusta population grouped with the *Coffea* strains from the pre-1970s outbreak (Flood 2005), suggesting it arose from a subset of older strains from the initial outbreak whereas the arabica population is more divergent. Thus, we first tested the hypothesis that the robusta population derived from the pre-1970s outbreak with the arabica population emerging separately. We then compared putative effector genes between the strains. Specifically, we ask (i) whether the 1990-2000s robusta epidemic is genetically different to the earlier outbreak; (ii) whether the host-specialist robusta and arabica populations share similar sets of derived effector genes or whether their similar pathologies evolved independently; and (iii) we test whether changes in pathogenicity and host-specialism involved horizontal transfer of effector genes and mobile chromosomes, or was restricted to within-lineage evolution in ancestral sets of effector genes using comparisons with potentially co-occurring and closely related *Fusarium* species.

## 3 Results

### 3.1 General features of the genomes in comparison with other *Fusarium* species

We reconstructed high-quality genomes ranging in size from 58 Mb in the *Coffea* strains to 61 Mb for the robusta strains and 63 Mb for the arabica strains. All had high contiguity (N50 ¿ 40 kb) and 100% genome completeness based on the presence of BUSCO genes (table 1). To evaluate our genomes further, we compared them to published genomes from a range of *Fusarium* taxa (table S1, figures 2, S1 and S2). *F. xylarioides* has a larger genome than its closely related species from the Gibberella fujikuroi Complex (GFC) African clade (Kvas et al. 2009), *F. udum* (56.4 Mb, Srivastava et al. (2018)), which causes wilt on pigeon pea, and that of *F. verticillioides* (41.7 Mb, Ma et al. (2010)), which is a non-wilt plant pathogen of maize. The genomes are similar in size, however, to the more distantly related *F. oxysporum* f. sp. *lycopersici (Fol)* strain 4287 (60 Mb, AAXH00000000.1, Ma et al. (2010)). *F. xylarioides* scaffolds broadly matched the 11 syntenic core chromosomes shared by *F. verticillioides, F. oxysporum* and more distantly-related *Fusarium* taxa (Ma et al. 2010) (figures 2, S1 and S2). We therefore used reference-guided scaffolding to combine our scaffolds into these 11 putative core chromosomes, with the rest of the genome remaining in shorter un-aligned scaffolds (figure 3). We classified these shorter scaffolds into four groups based on the degree to which they are shared in other GFC species: those matching short un-aligned scaffolds in the *F. verticillioides* genome assembly (FV), those with no match in *F. verticillioides* but matching *F. udum* and other *F. xylarioides* strains (FXU, 12% of the total genome, table 1), those specific to *F. xylarioides* as a whole (FXS, just 0.2% of the genome), and lineage-specific scaffolds unique to each *F. xylarioides* strain (LS, 3.5% of the total genome). Genome size differences are largely explained by the FXU, FXS and LS scaffolds (table 1).

**Figure 2:**
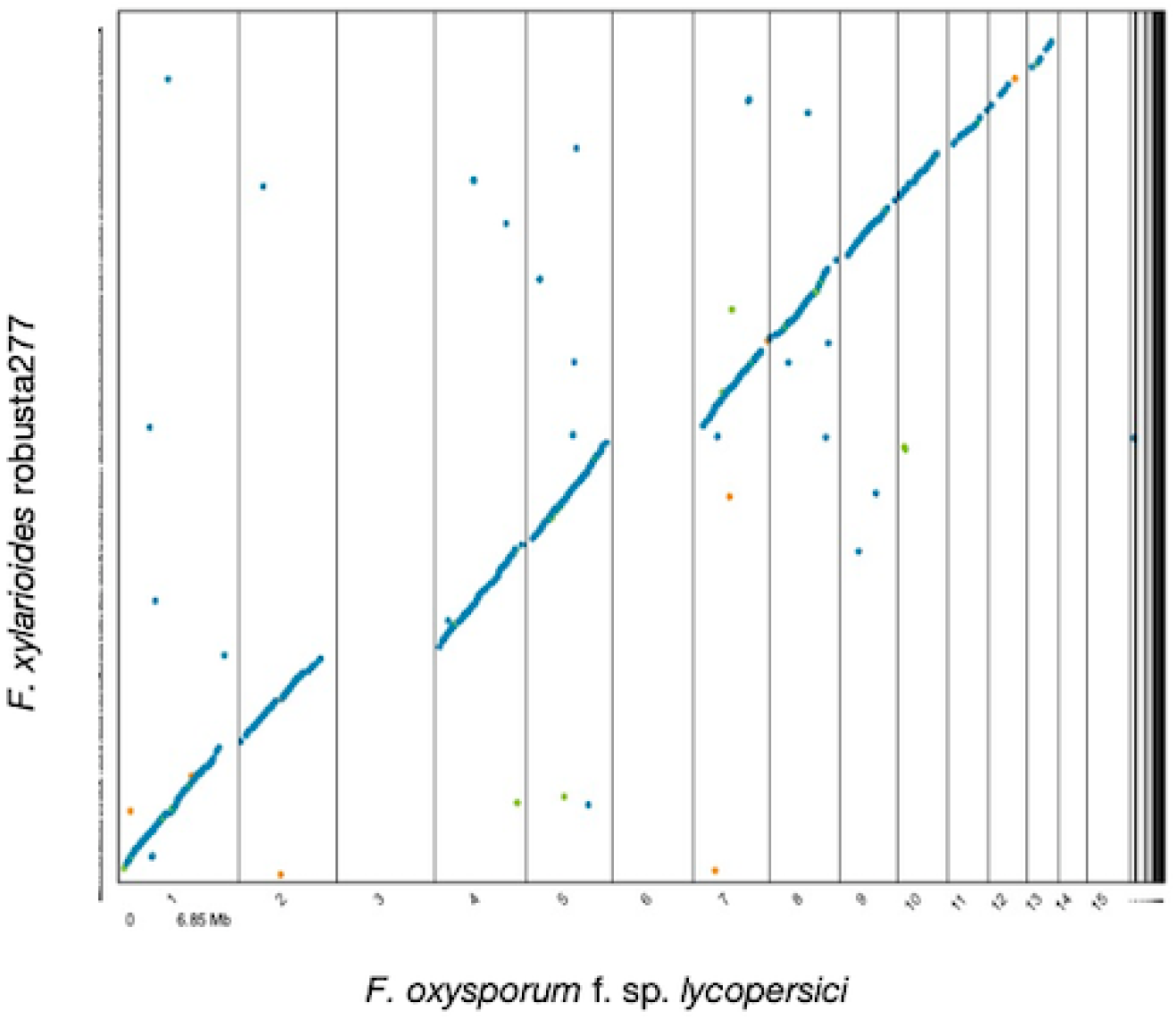
Representative whole-genome alignments of *F. xylarioides* strains against the 15 *F. oxysporum f. sp. lycopersici* chromosomes, including 11 core chromosomes shared with *F. verticillioides* and 4 mobile chromosomes (Ma et al. 2010). Each dot represents chromosomal correspondence, with absences representing the absent *Fol* chromosomes. Genomes were aligned using Mummer 4.0.0, with outputs processed using DotPrep.py before visualizing using Dot in DNANexus. Blue indicates forward alignments, green indicates reverse alignments, orange indicates repetitive alignments.

**Figure 3:**
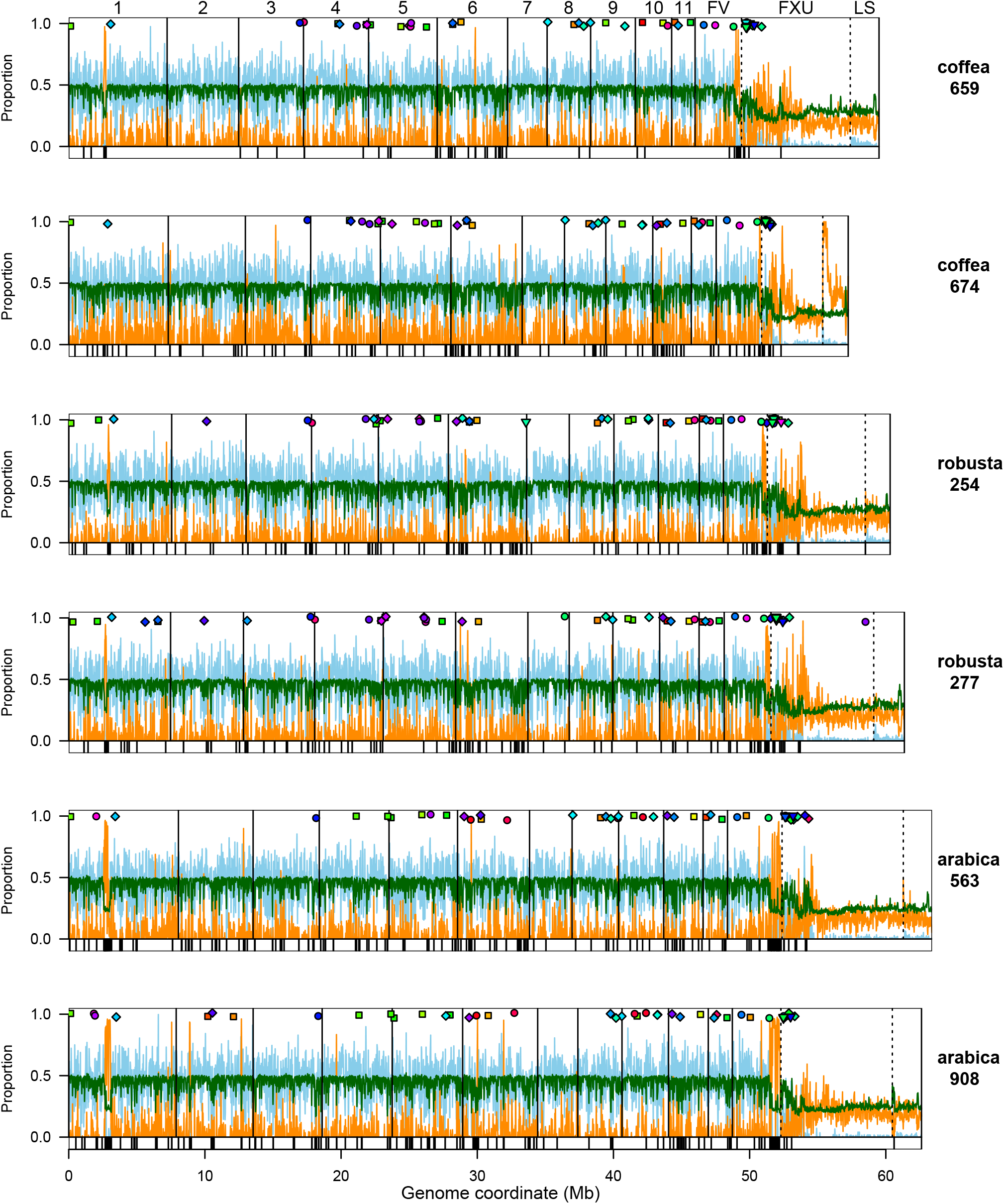
Global view of genome metrics plotted across 20 Kb windows of each *F. xylarioides* genome. Light blue = fraction of nucleotides annotated as a gene. Orange = fraction of nucleotides annotated as transposable element. Dark green = %GC. Vertical solid lines demark the 11 inferred chromosomes matching to the *F. verticillioides* genome. Dashed lines demark scaffolds that are shared with *F. verticillioides* but not assembled into chromosomes; shared with *Coffea659* ; or lineage specific. Coloured points indicate the location of putative effector genes: square = pre-defined in the literature, circle = CAZyme, diamond = small cysteine-rich proteins. Colours allocated across spectrum arbitrarily but same colour indicates belongs to same ortholog. The lower strip on each plot shows Large RIP Affected Regions in black

**Table 1:** Genome statistics for sequenced *F. xylarioides* strains compared with sister species. Abbreviations for *Fusarium* sister species: *Fu, F. udum; Fol, F. oxysporum f. sp. lycopersici; Fv, F. verticillioides*

| Strain number | 392674 | 127659i | 392277 | 392254 | 389563 | 375908i | *Fu* * | *Fol* * | *Fv* * |
| --- | --- | --- | --- | --- | --- | --- | --- | --- | --- |
| Short name | Coffea674 | Coffea659 | robusta277 | robusta254 | arabica563 | arabica908 |  |  |  |
| Other numbers | CBS258.52 | IMI127629 |  |  |  |  |  |  |  |
|  |  | DSMZ62457 |  |  |  |  |  |  |  |
| Population | Coffea | Coffea | Robusta | Robusta | Arabica | Arabica |  |  |  |
| Date isolated | 1951 | 1955 | 2003 | 1997 | 2002 | 1997 |  |  |  |
| Origin | Cote d'Ivoire | CAR | Tanzania | Uganda | Ethiopia | Ethiopia |  |  |  |
| Host | Coffea | C. excelsa | C. c. robusta | C. c. robusta | C. arabica | C. arabica |  |  |  |
| Genome Size (Mb) | 57.2 | 59.6 | 61.3 | 60.3 | 63.4 | 62.6 | 56.4 | 59.9 | 41.7 |
| No. scaffolds | 3969 | 4843 | 5484 | 5369 | 7,078 | 7,264 |  |  |  |
| N50 (bp) | 62845 | 55667 | 54978 | 56,291 | 42,705 | 41,762 |  |  |  |
| Chromosomes (Mb) | 51 | 49 | 51.6 | 51.3 | 52.4 | 52.3 |  |  |  |
| BUSCO score % | 100 | 100 | 100 | 100 | 100 | 100 |  |  |  |
| GC content % | 43.3 | 43.5 | 43.1 | 43.1 | 42.1 | 42.2 |  |  |  |
| Coding genes | 14,529 | 14,732 | 14,852 | 14,588 | 14,589 | 14,654 | 14,919 |  |  |
| Coverage % | 36 | 35 | 34 | 34 | 33 | 33 |  |  |  |
| Total repeats (Mb) | 13.3 | 14.9 | 15.9 | 15.3 | 18.6 | 17.8 | 12.6 | 10.8 | 1.0 |
| Total repeats (%) | 23 | 25 | 26 | 25 | 30 | 29 | 22 | 18 | 2 |
| Transposons (Mb) | 5.8 | 4.9 | 5.8 | 4.3 | 5.0 | 5.4 | 18.9 | 7.0 | 0.5 |
| Transposons (%) | 10 | 8 | 9 | 7 | 8 | 9 | 33 | 12 | 1 |
| FXU scaffolds (Mb) | 4.5 | 7.9 | 7.5 | 7.2 | 8.9 | 7.7 |  |  |  |
| FXS scaffolds (Mb) | 0.04 | - | 0.06 | 0.04 | 0.04 | 0.5 |  |  |  |
| LS scaffolds (Mb) | 1.9 | 2.2 | 2.3 | 1.8 | 2.1 | 2.1 |  |  |  |
| FXU scaffolds % | 8 | 13 | 12 | 12 | 14 | 12 |  |  |  |
| FXS scaffolds % | 0.1 | - | 0.1 | 0.1 | 0.1 | 0.8 |  |  |  |
| LS scaffolds % | 3 | 4 | 4 | 3 | 3 | 3 |  |  |  |
| FXU TEs (Mb) | 1.2 | 1.9 | 1.8 | 1.4 | 1.6 | 1.7 |  |  |  |
| LS TEs (Mb) | 1.0 | 0.6 | 0.6 | 0.4 | 0.5 | 0.4 |  |  |  |
| Chromosome |  |  |  |  |  |  |  |  |  |
| TEs (Mb) | 3.6 | 2.4 | 2.8 | 2.4 | 2.9 | 3.2 |  |  |  |
| FXU TEs (%) | 28 | 24 | 23 | 20.1 | 18 | 22 |  |  |  |
| LS TEs (%) | 52 | 26 | 28 | 24 | 23 | 20 |  |  |  |
| Chromosome |  |  |  |  |  |  |  |  |  |
| transposons (%) | 7 | 5 | 5 | 5 | 6 | 6 |  |  |  |

*F. xylarioides* and *F. udum* are the only members of the GFC to induce wilts (Rutherford 2006, Geiser et al. 2005). To test whether they might have evolved new wilt capabilities by acquiring a pathogenic chromosome from *Fol*, we mapped the *F. xylarioides* genomes to the *Fol* genome assembly. Chromosomes 3, 6, 14 and 15 in the *Fol* genome are supernumerary, or mobile, as well as parts of chromosomes 1 and 2 (Ma et al. 2010). Chromosome 14 is also pathogenic, housing all known *Fol* wilt effector genes. We found no large-scale matches to any of the fully mobile chromosomes in *F. xylarioides* (figures 2 and S2, ruling out the transfer of the whole *Fol* mobile pathogenic chromosome. However, there was a significant excess of smaller matches between LS and FXU scaffolds and the *Fol* mobile chromosomes (figure S3, Pearson’s chi-sq test: p¡0.001), consistent with smaller pieces having been transferred: 7% of the FXU scaffolds and 20% of the LS scaffolds matched to the four fully mobile *Fol* chromosomes (3, 6, 14 and 15), whereas <2% matched to non-mobile chromosomes. It is possible therefore that acquisition of wilt-formation in the GFC could have involved transfer of smaller genome regions or individual pathogenicity genes from *Fol* or a related *F. oxysporum*.

Annotation of genes and transposable elements (TEs) shows that the scaffold groups in *F. xylarioides* differ considerably in overall genomic content. The 11 core chromosomes are gene rich and repeat poor, whereas the FXU, FXS and LS scaffolds are gene poor and repeat rich (figure 3). The whole genomes contain on average 14,650 predicted protein-coding genes, covering approximately 34% of the genome (table 1). The distribution of genes attributed to particular functional classes was highly concordant among the six *F. xylarioides* strains and *F. udum* (figure S4), and among every core chromosome except chromosome 11, which is enriched for putative functions related to stress response and protection from reactive oxygen species (figure S5). In contrast, the FXU, FXS and LS scaffolds are enriched for transposable elements, which overall comprise 24% of DNA in these scaffold groups, compared to 5% in the 11 core chromosomes. Within this, retrotransposons have contributed to genome expansion in *F. xylarioides*, as is commonly observed in other fungi (Schmidt et al. 2013), making up 16% of DNA in these scaffold groups, compared to 3% in the 11 core chromosomes. The LS scaffolds are further enriched for DNA transposons, with 150% more than is found in the FXU scaffolds and 360% more than in the core chromosomes. Interestingly, genes on FXU and LS scaffolds are also enriched for the putative function ‘phage major capsid proteins’ (figure S5), which could be involved with movements of mobile elements. The proportion of the whole genome made up of transposable elements also varies among species. In line with the idea that transposable elements provide a genetically variable environment within which new pathogenicity traits can develop, the three wilt-inducing species *F. xylarioides*, *F. udum* and *F. oxysporum* all have higher repetitive sequence and TE content than non-wilt formers. For example, *F. xylarioides*, has nearly 1000% more TEs than *F. verticillioides* (tables 1 and S2), consisting of on average 16 Mb of total interspersed repeats with nearly one third of these (5.2 Mb) recognised as retroelement and DNA transposons. In *F. xylarioides*, the remaining two thirds are unclassified, while *F. udum* and *Fol* have much lower proportions of unclassified repeats (between 30-40%) (table S2).

The *F. xylarioides* genomes display another common structural feature of plant pathogenic ascomycetes, namely the presence of AT-rich genome blocks. Specifically, the Repeat-Induced Point-like (RIP) mutations pathway is a genome defence mechanism against the invasion of mobile transposons that has been linked to enhanced mutagenesis of effector genes in some plant pathogens (de Jonge et al. 2011). It is unique to ascomycete fungi and mutates cytosine bases to thymine in repeated genome sequences (Van Wyk et al. 2019). This leads to AT-rich sequences that are typically low in coding gene content, and can segment the genome into equilibrated compartments (as in *Leptosphaeria maculans*, (Rouxel et al. 2011)). We find significant evidence for AT-rich blocks (averaging 21.5% GC), which are spread throughout the core chromosomes, and the FXU and LS scaffolds (figure 3). On average, these blocks are significantly poor in genes but rich in repeats, in common with patterns in other fungi. We find that RIP regions make up >33.5% to >40% of each genome, with >7.7% to >8.5% of the genomes in so-called Large RIP Affected Regions (LRARs, (figure 3). Having described the broad features of our genomes we now address our questions concerning the multiple outbreaks.

### 3.2 The arabica population arose independently from the robusta and *Coffea* strains

Our genome data supports the previous evidence that the arabica and robusta populations emerged independently within *F. xylarioides* (Flood 2005). Gene annotations from the *F. xylarioides* and *F. udum* strains together with ten other published *Fusarium* and *Verticillium* wilt genomes were used to identify 18 569 orthologous gene sets, encompassing 25,0056 genes or 95.6% of all annotated genes. The species tree based on the concordance of gene trees of all ortholog groups supports the established order of the GFC (Kvas et al. 2009) as well as the monophyly of *F. xylarioides*, with over 87% of genes supporting monophyly of the clade (figure 4). No alternate topology was consistently found for the remaining genes trees (the next most common was supported by just 1.1

**Figure 4:**
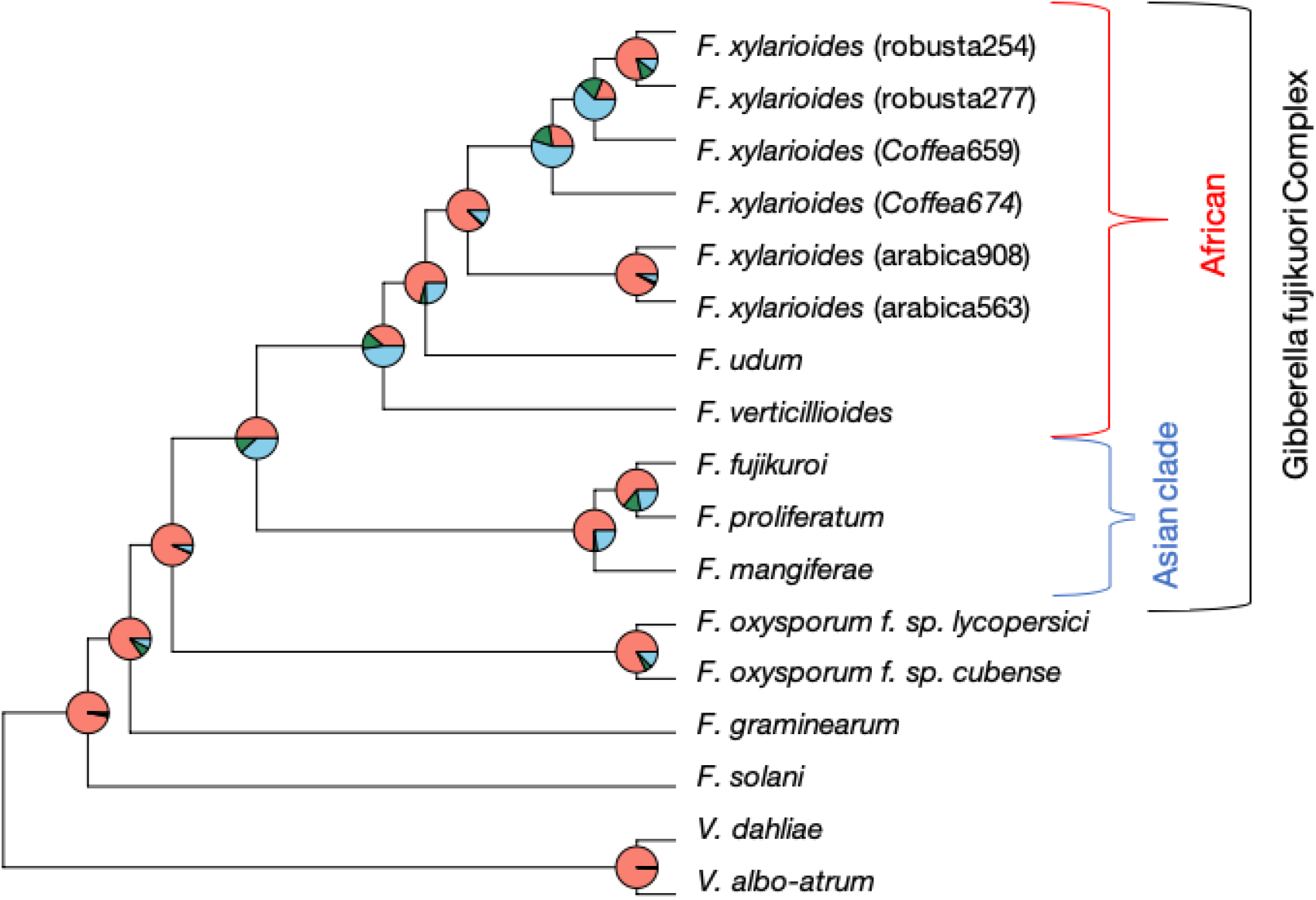
Phylogenetic relationships between *Fusarium* species reconstructed from 13 782 orthogroups support monophyly of the *F. xylarioides* clade, with little consistent support for alternate topologies. Strain accession numbers are in table S1. The brackets denote the GFC, with the Asian (blue) and African (red) clades. Pie chart colours: pink = proportion of genes (orthogroups) recovering the depicted node; dark green = the proportion of genes recovering the second most common topology; light blue = the proportion of genes recovering all other topologies.

This conclusion is further supported by patterns of presence and absence of genes. Over 95% of single gene copies were shared between the two arabica and between the two robusta strains respectively, while there is less similarity between other *F. xylarioides* comparisons (arabica-robusta, *Coffea-Coffea*, and *Coffea*-arabica/robusta, figure 5a). The robusta strains share a significantly higher proportion of orthologous gene sets with *Coffea659* (SuperExactTest, p¡0.001), and they generally display more concordance with the *Coffea* strains, while the arabica strains have the most unique orthogroups. The arabica strains share slightly more genes with the ex-type strain *Coffea674* than with *Coffea659* despite the latter being the only strain able to infect both both arabica and robusta coffee (Girma et al. 2009, Flood 2005).

**Figure 5:**
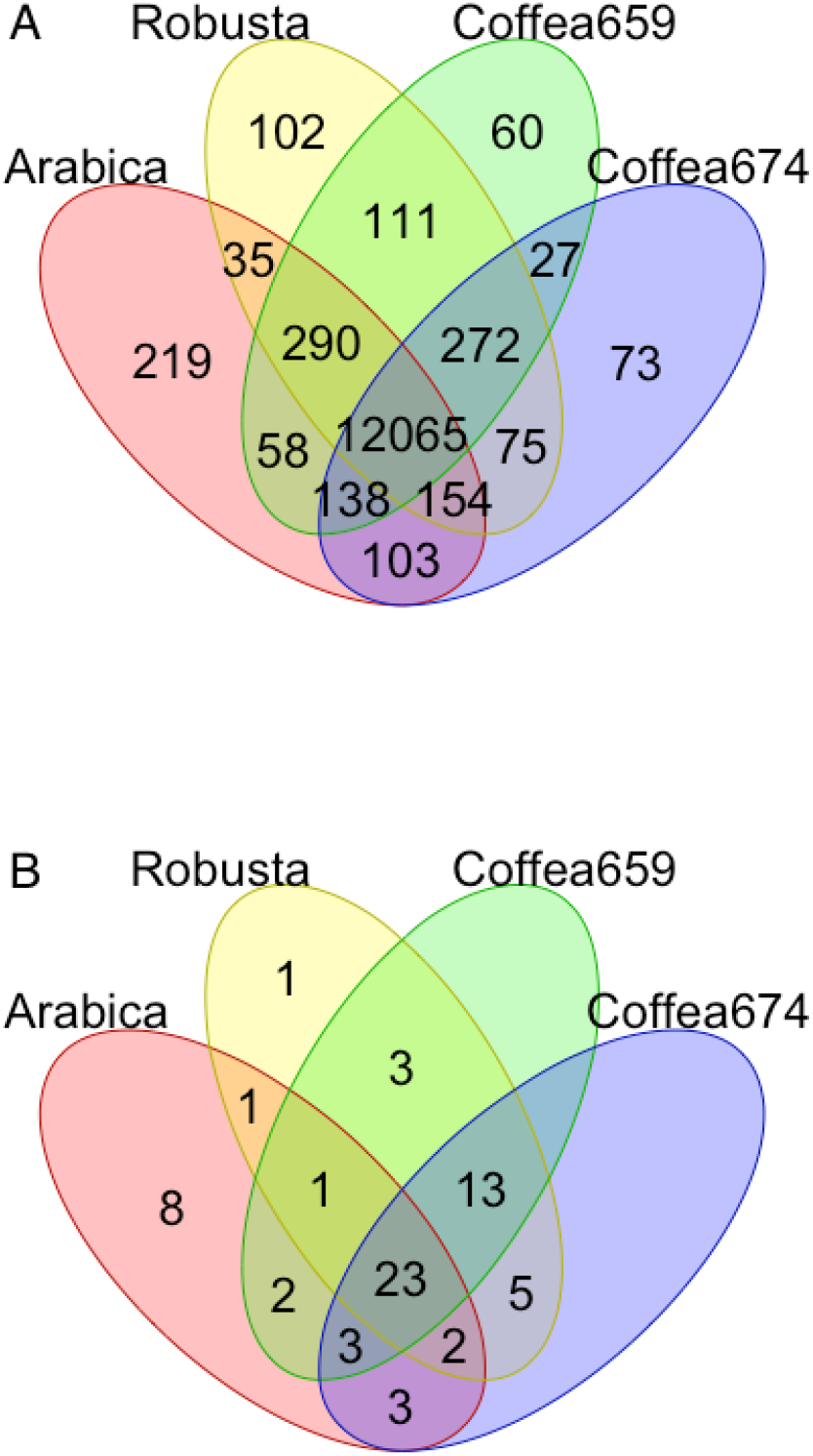
A) Orthgroups shared between *F. xylarioides* arabica, robusta and the two *Coffea* strains. Drawn using 13 782 orthogroups (excluding 449 *F. xylarioides* robusta orthogroups and 175 *F. xylarioides* arabica orthogroups that differed in presence/absence between the the two strains of each population. The two *Coffea* strains are more divergent (with 1123 orthogroups that differ in presence/absence between the two strains) and so are drawn separately. The two arabica strains (563 and 908) share 13 062 genes and the two robusta strains (254 and 277) share 13 104 genes in total. B) Putative effectors shared between *F. xylarioides* arabica, robusta and the two *Coffea* strains. Drawn using 66 putative effector proteins that differed in presence/ absence across the host-specific populations.

### 3.3 A set of putative effector proteins for *F. xylarioides*

The proteome of plant pathogens includes pathogenicity factors and effector proteins, which are secreted to enable pathogen entry, survival and effective colonisation in their host. Fungal effector proteins are typically under positive selection and do not show conserved protein sequences (Jones et al. 2018, de Jonge et al. 2011). Therefore, we adopted a multi-dimensional approach to look for putative effector proteins, namely those which enable the pathogen to become established in its host and trigger the onset of symptoms and a host defence response. We described three classes: *(i)* those known from other fungal pathogens, *(ii)* small and cysteine-rich proteins, and *(iii)* carbohydrate-active enzymes. Altogether, this process identified 65 putative effector genes, which were distributed across the 11 core chromosome scaffolds as well as the LS and FXU-specific scaffolds (figure 3), and mostly recovered in the same location in each strain (figure 6). Because effector genes have not been investigated previously in *F. xylarioides*, we first describe general findings for each category of effector genes, before focusing on the differences between the different host-specific populations.

**Figure 6:**
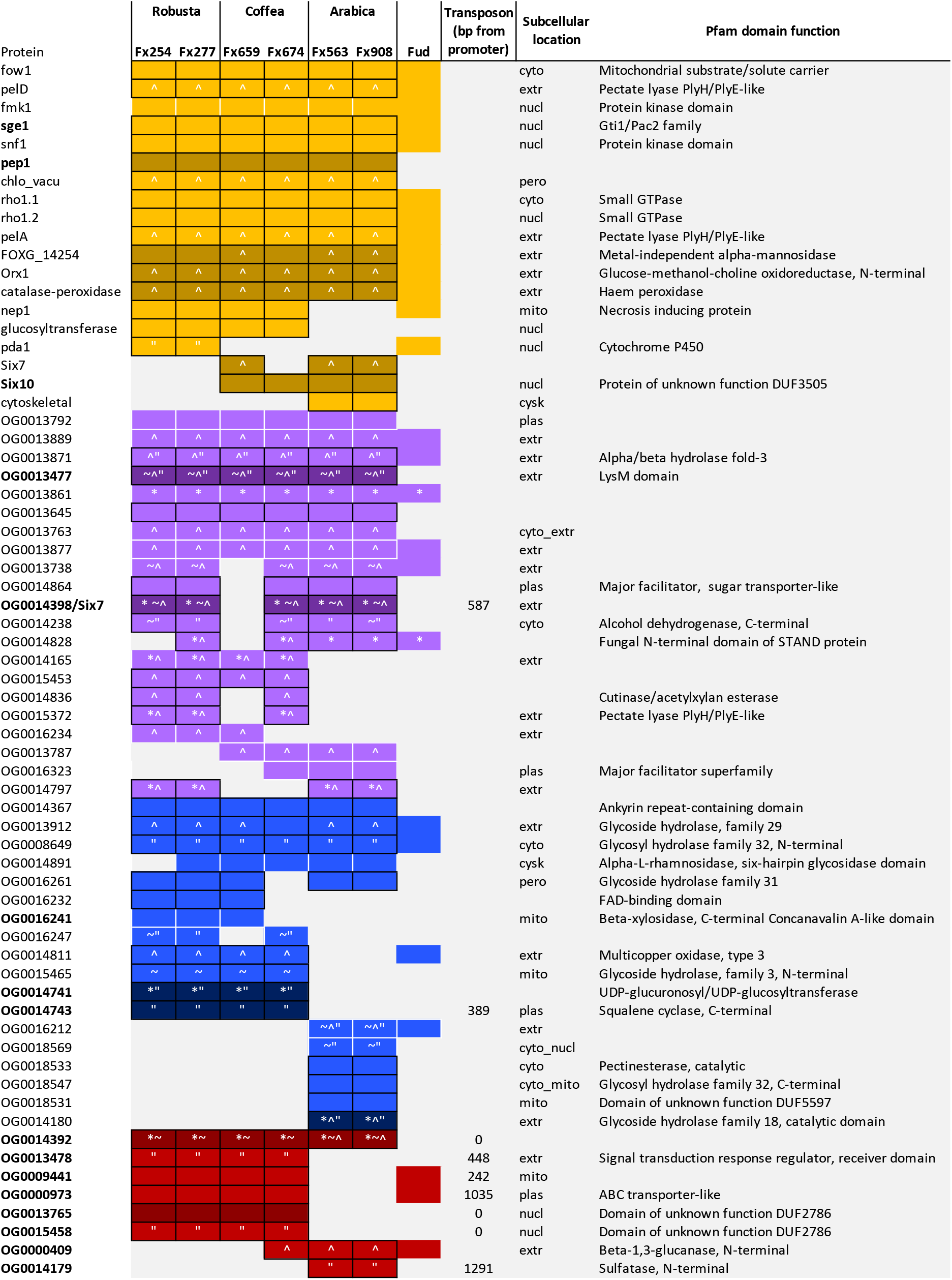
Putative effectors’ characteristics and presence or absence across *F. xylarioides* strain and *F. udum*. The four effector classes are shown in: yellow for predefined effectors; purple for small and cysteine-rich secreted effectors; blue for carbohydrate-active enzymes; and red for transposon-adjacent effectors. The presence of transposons is represented by names in bold with its distance from the genes promoter described if less than 1500bp (if not, the transposon is over 1500bp away), genes under positive selection by an asterisk, genes in an AT-rich region by a tilde, genes which share a locus across all strains are outlined in black, genes with evidence of horizontal transfer from *F. oxysporum* are a darker shade and function is represented by the Pfam domain, where a hit was returned.

#### 3.3.1 Putative effector genes known from other species

We found BLAST matches in *F. xylarioides* to 18 known effector proteins previously characterised in *Fol* and other wilt-inducing pathogenic fungi, including some that are only recently discovered and unnamed (table S3, figure 6). The effector genes *fow1, fmk1, snf1, pelD, chlo-vacu, sge1, rho1* (here with two copies *rho1.1* and *rho1.2*), *pep1, pelA, orx1* and *catalase-peroxidase* were found in all six *F. xylarioides* strains, with *nep1, gluco, pda1, six7, six10, cytoskeletal* found across the host-specific pairs. The majority of the well-characterised SIX effectors from *Fol* chromosome 14 are absent in both *F. udum* and all *F. xylarioides* strains, with the exceptions of the recently described Six10 protein (Schmidt et al. 2013) which is present in arabica and *Coffea* strains, Six7 is present in arabica and *Coffea659*, and the transcription factor Sge1, required for the expression of SIX genes (Michielse & Rep 2009), is found in all strains.

#### 3.3.2 Small and cysteine-rich secreted proteins

Many fungal effector proteins are small (¡400 amino acids), cysteine-rich (¿4 cysteine residues) and secreted (Van Esse et al. 2008, de Jonge et al. 2011). Searching our annotated proteomes for these properties discovered over 2,500 small, cysteine-rich proteins in each strain, of which over 500 also have a signal peptide and a signal peptide cleavage site. Focusing on single copy genes for ease of comparison left 132 putative effectors in 21 orthologous groups shared across *Fusarium* and *Verticillium* (figure 6). Patterns of presence and absence were variable among these genes: only 5 were recovered in all *F. xylarioides* strains and *F. udum*. Many of these genes had no recognisable Pfam domain functions identified by InterProScan (figure 6). This is to be expected because there are large numbers of unknown small and cysteine-rich proteins secreted in the apoplast (as part of the water-transport pathway) which are typically species- or strain-specific which have no known function (de Jonge et al. 2011). However, two orthogroups (OG0013477 and OG0013645, figure 6) had BLAST matches to the LysM-containing chitin oligosaccharide protein, which in *Cladosporum fulvum* is required for scavenging of chitin oligosaccharides as a conserved fungal defence to enable undetected invasion (De Jonge et al. 2010). Orthogroup OG0014398 displayed a match, albeit with a relatively low BLAST score, to a number of *six7 F. oxysporum* genes and protein sequences (notably not the *Fol* 4287 sequenced genome used for this comparative analysis) (figure S6), which might indicate that it is a divergent type of six7 effector compared to previously described copies. Other functional annotations reveal a recurring theme of pectin degradation (expanded in next section): OG0013871 is a hydrolase; OG0015372 is a pectin lyase and OG0014836 a carbohydrate esterase 5 with a carbohydrate binding module (CBM1); and OG0014864 is a major facilitator superfamily involved in the transport of solutes, which could influence wilting.

#### 3.3.3 Carbohydrate-active enzymes

Carbohydrates in plant cell walls such as cellulose and pectin provide the main source of carbon for fungal pathogens (Hervé et al. 2010) and so carbohydrate-active enzymes (CAZymes) are thought to be important in the infection pathway. We therefore searched for CAZymes restricted to some or all of the wilt-inducing species of *Fusarium* and *Verticillium* and classified their carbohydrate-binding modules (CBMs) to characterise specificity for particular polymers. Our search found 60 genes of interest belonging to 19 orthogroups (figure 6). As a whole, *Fusarium* is broadly enriched for lytic enzymes involved in carbohydrate metabolism along with other ascomycete fungi (Reis et al. 2005, Soanes et al. 2008) (table S4), while the vascular wilt inducers show enrichment in certain gene families (table S5), also reported by Klosterman et al. (2011)). Fungal growth on pectin requires either the combined activities of pectin esterases and polygalacturonases or the sole activity of pectate lyases (Howell et al. 1976, Mace et al. 1981), and all three enzymes are known from other wilt-inducing fungi (Beckman 1956, Langcake & Drysdale 1975, Talboys & Society 1970). Interestingly, each share orthologus groups across the wilt-inducing strains analysed in this study (table S5). A polygalacturonase enzyme (glycoside hydrolase (GH) family 28) from *F. oxysporum*, has expanded by one copy number in all *F. xylarioides* strains except *Coffea674*. Pectate lyase carbohydrate-binding modules CBM66 and CBM38, which bind a fructose-hydrolysing GH32, and chitin-cleaving CBM18 and CBM50 are also found across all six *F. xylarioides* strains (as well as *F. udum* and *Fol*). In contrast pectin esterases have expanded differentially among different *F. xylarioides* strains (see next section).

### 3.4 Host-specific populations differ in their complement of putative effectors

Effector gene presence and absence recapitulated findings from the whole genome analyses: both strains from the same host-specific population nearly always share the same effector gene complement, and the robusta population and *Coffea* strains share greater overlap in effector gene complement (SuperExactTest, p=0.001) than either do with the arabica population (figure 5b). Based on this significant association, we looked for changes that might have led to specialisation on robusta coffee and increased virulence, and found just 1 effector unique to the robusta strains, *pda1*. This gene encodes the enzyme pisatin demethylase, which has been shown to detoxify the plant defensive compound (phytoalexin) pisatin by *F. oxysporum f. sp. pisi* in peas, and with different alleles present across different *F. oxysporum* host-specific f. sp.’s (Covey et al. 2014, Ellis et al. 2016, Coleman et al. 2011). Interestingly, in sugarbeet one *pda1-a* allele (accession AY487143.1) was only found in pathogenic strains (Covey et al. 2014), although the gene copy found in this study is 97% similar to a *pda1* from *F. oypsorum f. sp. phaseoli* and is only 62% similar to this pathogenic sugarbeet allele. The enzyme could have a direct role against related phytoalexin compounds in coffee or enable growth of robusta strains on alternative hosts. However, with the excpetion of *pda1*, the robusta population is more unique in genes that it lacks than in genes it has gained. Because plant immune systems are triggered by detection of secreted effector proteins, the loss of genes might also be important for pathogenicity on different hosts, should plants evolve to recognise certain effectors. Four orthogroups that are present in one or both *Coffea* strains are missing from the robusta strains: *six7, six10, OG0013787* and *OG0016323* so absence of these genes could be further explored as a possible cause of enhanced pathogenicity on robusta.

In contrast, the arabica population is highly divergent to both the *Coffea* strains and the robusta population in its complement of effector genes. Arabica strains share 7 unique effectors: *cytoskeletal* and 6 CAZymes. The wilt-specific Cytoskeletal protein was recently described in *Fol, V. dahliae* and *V. albo-atrum* and found to be absent from non-wilt inducing *Fusarium* species (Klosterman et al. 2011). Among the CAZymes, *OG0018533* is a pectin esterase, and *OG0014180* encodes for a GH18 chitinase that has an insertion under positive selection relative to the matching copy in the *Fol* genome, potentially indicating divergent function within the arabica population. In addition, the arabica strains lack 15 effector orthologs found in one or both *Coffea* strains: *gluco, nep1*, 5 small cysteine-rich proteins including a hydrolase and a pectate lyase, and 8 CAZymes including 3 more hydrolases and another glucosyltransferase, *OG0014741*. These two glucosyltransferases are found across the robusta and *Coffea* strains, *F. oxysporum* f. sp.’s including *Foc, Fol*, *V. dahliae* and *V. albo-atrum*. The predefined *gluco* glucosyltransferase is required for full pathogenicity in *V. dahliae* Klosterman et al. (2011), and in *F. xylarioides* shares a scaffold with the small cysteine-rich putative effector *OG0014165*, which is also only present in the robusta and *Coffea* strains (see below). The Nep1 protein induces necrosis and ethylene production in host plants (Pemberton & Salmond 2004) with its family expanded in *Fol* and *V. dahliae* and purportedly responsible for their broad host ranges (Yadeta & J. Thomma 2013). It is noteworthy that no putative CAZyme effectors are shared exclusively between arabica and the *Coffea* strains, whereas the robusta strains share seven with the *Coffea* strains.

While these differences confirm separation of the arabica population from the other strains, a few putative effectors displayed contrasting affinities. The four genes highlighted as absent in robusta strains are cases of sharing between *Coffea* and arabica strains. Of these, *six7* is shared with *Coffea659*, the only *Coffea* strain that is also able to infect arabica coffee (Girma et al. 2009, Flood 2005), and therefore of possible interest as pathogenicity factors for growth on arabica coffee. Just one gene, *OG0014797*, is shared by robusta and arabica populations but absent in both *Coffea* strains. Interestingly, this gene gave a significant signal for positive selection and displays considerable amino acid divergence between the two forms. However, both copies share *F. anthophilum*, a member of the American GFC clade (Kvas et al. 2009), as closest relative (with nearly 80% similarity), while a diverged copy in *F. verticillioides* is closer to *F. nygamai* (84% similarity), another African GFC clade member (Kvas et al. 2009). It is possible therefore that this gene diverged rapidly under positive selection for differential function in each population or that these represent separate acquisitions from alternative sources, despite the same closest BLAST match for *F. xylarioides*. Either way, there is no evidence for substantial recent exchange of effector genes between arabica and the other populations, consistent with genome-wide evidence for their isolation.

We also looked for changes in the number of copies of CAZyme gene families shared by orthologous groups among *F. xylarioides* strains as a possible cause of changes in the ability to degrade plant cell walls and access carbon from their host. Excluding the groups which matched with non-wilt inducing species, we found differences in vascular wilt-inducing pectin-degrading gene families between robusta and arabica populations, with the *Coffea* strains, as before, more similar to robusta. Pectin esterases in the carbohydrate esterase family 8 (CE8) and the pectate lyase PL1 4 have expanded in the arabica population only, whereas the robusta population has expanded in the CE5 family and a CBM35 gene, which both hydrolyse xylan instead, a component of hemicellulose also used for carbon by wilt fungi (Lombard et al. 2013, Mace et al. 1981). Both CE5 and CE8 families are also present in *Fol* and *Verticillium*. Whilst not specific to pectin, CBM1-containing carbohydrate esterases that bind cellulose and hydrolyse xylans, mannans and pectins (Kellett et al. 1990, Hogg et al. 2003, McKie et al. 2001) were only present in robusta strains and *Coffea674*. While we cannot infer the functional effects of these changes on the infection process, they are suggestive of differences in carbon usage between the two host-specific populations.

Although the main differences among strains are apparent in gene content, we also found cases of divergent selection acting on the amino acid sequence of some shared genes: for example, OG0013871, a Carbohydrate-Binding Module 9 (CBM9 xylanase) and OG0014398, found in all strains except *Coffea659* (figure 6. The latter ortholog is the potentially divergent form of *six7* identified as a small cysteine-rich protein above and displays amino acid divergence in a pattern consistent with strain host-specificity (figure 7).

**Figure 7:**
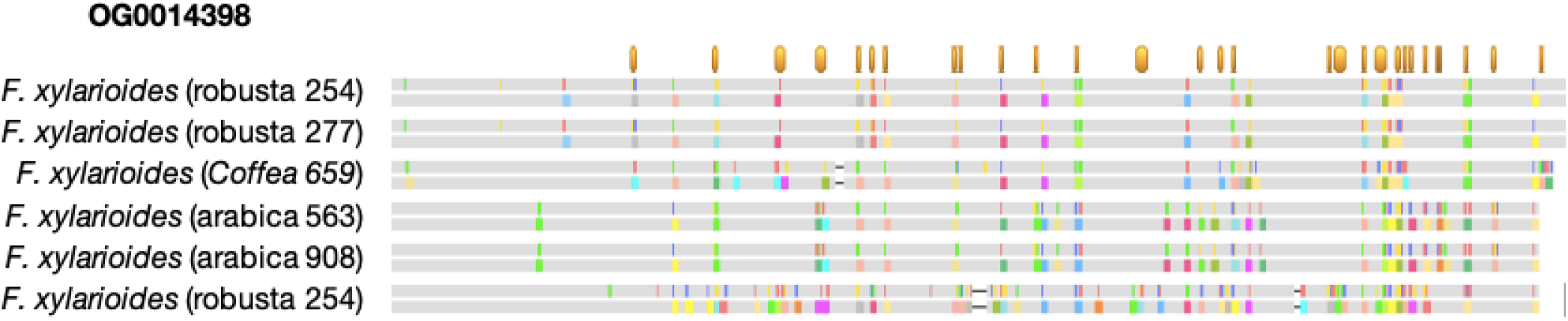
The translated protein alignment of OG0014398. No changes between nucleotide (top grey bar for each taxon) and translated protein (bottom grey bar) between the strains are shown in grey, with changes to either marked in colour. Codons under positive selection (dN/dS ratio > 1 in PAML Bayes Empirical Bayes analysis) are noted by an orange bar along the top. Robusta254 has two gene copies in this orthologous group and Coffea674 has none. Drawn in Geneious 9.1

### 3.5 Several effector genes have been acquired horizontally by transposable elements

Despite there being no evidence for transfer of whole known mobile chromosomes from *Fol*, we do find evidence for the acquisition of specific effector proteins associated with mobile elements. Twelve putative effectors have close BLAST matches to *Fol* mobile chromosomes. Of these, three match to mobile chromosomes that were not previously linked with pathogenicity: *six10* and *pep1* match regions of chromosomes 3, 6 and 15, while *OG0014180* matches 3 and 6. The remaining nine effectors have a BLAST match to *Fol* chromosome 14 (*OG0014398, OG0013477, OG0014741, OG0014743, OG0013765*, *six7, FOXG 14254, orx1, catalase-peroxidase*) with the first six additionally absent from other closely-related *Fusarium* species (figure 6). Intriguingly, eight of these effectors all match regions within chromosomal mini-clusters on chromosome 14 that contain *Fol*’s SIX effectors (being within 1.8kb to 33kb of SIX genes, figure S7, Schmidt et al. (2013)). These mini-clusters are enriched in class two DNA transposons and have miniature Impala (mimp) transposons adjacent to their promoter regions (van Dam & Rep 2017, Schmidt et al. 2013). Schmidt et al. (2013) suggest that effector genes found between interspersed repeat regions on chromosome 14 could coincidentally be transposed with the mimp transposons which could explain the presence of these effector genes in *F. xylarioides*.

Consequently, we looked for further cases of effectors that are close to transposons. In addition to mimps, Schmidt et al. (2013) further described several new transposons in *Fol* that might promote transposition or mutation of effector genes: a class 1 retrotransposons *yaret1* from the LINE (long-interspersed nuclear elements) superfamily and *yaret2* from the LTR (long terminal repeat) superfamily, which encodes a retroviral integrase that inserts a DNA copy into the host genome; and a class 2 DNA transposon *hop3* from the Mutator family. We therefore investigated the distribution of these elements in our genomes. The LS scaffolds house on average 30% of all Impalas, and 75% of mimps, which is proportionally far more than are found on the core chromosomes (table S6). The robusta strains contain the most mimps and the arabica strains have the fewest. Next, we examined the promoter regions (defined as 1500 bp upstream from the start codon) of genes that met four criteria: they were present in both strains for each host-specific population; they contained mimps or the newly described transposon families from Schmidt et al. (2013); they were on FXU or LS scaffolds; and they matched regions of *Fol*’s mobile chromosomes. This resulted in eight additional putative effectors to supplement those identified by our three original criteria, with three containing mimps, two containing *hop3*, two containing *yaret2* and one with *yaret1* (figure 6). Additionally, eight putative effectors from the predefined, small and cysteine-rich and CAZyme classes also shared a scaffold with a transposon, with *OG0014398* (*six7*) and *OG0014743* with mimps adjacent to their promoter regions.

Among these putative effectors with evidence for acquisition by transposition, seven are found in both robusta and arabica populations and at least one *Coffea* strain. Of these, one is the putative divergent form of *six7* (*OG0014398*) that displays evidence of divergent selection between host-specific strains. Among the rest, we can judge the potential contribution of transfer from *F. oxysporum* to the emergence of the host-specific races. Three are shared by *Coffea* and robusta strains (OG0014741 a glucosyltransferase, OG0014743 a squalene-hopene-cylase, and a protein of unknown function), two are shared by *Coffea* and arabica strains (Six7 and Six10), and one is restricted to the arabica population (OG0014180, a GH18 chitinase). Thus, while there has been a notable contribution of transfer, for the majority of differences in putative effector gene content among host-populations we found no evidence of horizontal transfer from *F. oxysporum*. BLAST searches against Genbank found *F. oxysporum* to be the closest match to 39 putative effectors, with no other possible donor consistently found (figure S8). Of these, 25 are found in neither *F. graminearum* nor the Asian GFC clade species nor *F. verticillioides* (figure 4). Indeed, 14 putative effectors are absent from these species and share a very high sequence similarity with a 90% or higher match with *F. oxysporum* (figure S8), suggesting the transposition of these genes. Of the *formae speciales* which consistently match, *F. oxysporum f. sp. raphani* shares a percent identity >80% with four effectors only found in the robusta and *Coffea* strains and absent across sister *Fusarium* species. Of the further *F. oxysporum* strains named in matches, the *formae speciales pisi, vasinfectum, lycopersici* and *cubense* all match multiple effectors and are absent in sister *Fusarium* too.

Finally, effector loci are significantly associated with regions affected by RIP-mutation. Effector genes overlap significantly more with RIP regions than expected for the same number of randomly chosen genes. The proportion of total effector gene coding sequence overlapping RIP regions ranges from >47.0% to >60.2%, compared to upper >95% values of >38.9% to >47.7% for the same number of randomly chosen genes from the whole proteome (randomisation test with 100 trials, p¡0.05 in arabica908, p¡0.01 in all other strains). Those effectors that are specific to either robusta or arabica populations have a mean overlap with RIP regions of 100% and >47.3%, respectively. These results support the hypothesis that RIP-mutation provides an additional mechanism for enhancing variability of the effector genes.

## 4 Discussion

Using fungal strains collected over a period of 70 years and optimally cryopreserved in a living state, we uncovered the genomic basis of repeated outbreaks of *F. xylarioides* on different commercial coffee species. Our data support the conclusion that the robusta and arabica populations had divergent origins within the *F. xylarioides* clade, with the robusta population deriving from within the earlier outbreaks on alternative *Coffea* crops. Furthermore, by cataloguing multiple putative effector and pathogenicity genes, we show how the different host-specific populations acquired different genes by horizontal gene transfer from *F. oxysporum*, as well as diverging in gene content within lineages.

The finding that arabica is divergent from the other strains is consistent with earlier work based on crossing experiments and molecular markers, which suports the reclassification of the 1990-2000s host-specific populations as separate species named *F. abyssiniae* (arabica) and *F. congoensis* (robusta). Our evidence for concordant gene trees across the entire genome, as well as substantial differences in gene content, both overall and specifically for genes thought to be important for the infection process, lends weight to this proposal. Rather than the presumed recent emergence of these strains, the level of diversity we observe is consistent with a divergence far earlier than the emergence within the last 50 years of these populations as major disease agents. Moreover, we found little evidence for transfer of genes from the pre-1970s outbreak into the 1990-2000s epidemic on arabica coffee. We hypothesise that this represents a separate emergence on commercial arabica coffee from *F. xylarioides* strains associated with wild coffee relatives in Ethiopia or previously less noticeable disease symptoms on arabica, but further sampling including of wild relatives would be needed to test this. In contrast, the robusta population evolved with relatively minor modifications from the original outbreak on multiple *Coffea species* in central and west Africa, with only two putative effector genes gained and a handful lost compared with the *Coffea* strains sampled here. The high level of similarity between paired strains within each host-specific population, despite sampling from different locations or countries five years apart, supports the emergence of each population from low initial diversity, i.e. each constitutes a single epidemic outbreak replacing earlier variants (Flood 2005, Phiri & Baker 2009).

The observation that several effector genes share a high percent identity with *F. oxysporum* and display close matches to its pathogenicity chromosome, whilst lacking in other more closely related *Fusarium species*, supports the role of horizontal transfer in the origin of host-specific differences. The mobile pathogenic chromosome of *F. oxysporum* is widely reported to transfer pathogenicity between different strains (Ma et al. 2010, Schmidt et al. 2013). However, to our knowledge, the transfer of pathogenicity has not been reported across different *Fusarium* species, nor has it been linked with the other mobile chromosomes. Such transposition could have taken place in their shared niche, where both *F. xylarioides* and *F. oxysporum* (undescribed f. sp.) have been coisolated from the roots and wood of CWD-infected coffee in Ethiopia and central Africa, as well as from banana roots where banana and coffee were intercropped in Uganda (Serani et al. 2007, Flood 2005). *F. oxysporum* f. sp. *cubense* (*Foc*) is widespread on banana in east and central Africa (Serani et al. 2007). Alternately, given that there is evidence that *F. xylarioides* can infect tomatoes (Onesirosan and Fatunla, 1976) it is possible that the *Solanaceae* family is an alternate host and *Fol* is the source of pathogenicity. Other *formae speciales* which are the closest match to numerous effectors are *F. oxysporum* f. sp. *pisi* and *F. oxysporum* f. sp. *raphani* which could be investigated as additional possible sources of pathogenicity. If confirmed, this would suggest avoiding the widespread practice of intercropping species that share closely related pathogens, for example coffee and banana, or weed management for those which share pathogens, for example of *Solanaceous* weeds in crop surroundings. Greater sampling from these taxa and around coffee fields would be needed to further clarify the range of potential donors for horizontally acquired genes.

Contrary to previously described cases in *F. oxysporum* (Ma et al. 2010, Schmidt et al. 2013), we find no evidence that transfer occurred on a known mobile chromosome, and indeed many of our putative effector genes are spread across our genome scaffolds. It remains possible that an unknown mobile chromosome or chromosomes with some homology to the *Fol* mobile chromosomes is involved with some of the variation among strains that we observe: we were unable to assemble LS and FXU regions into whole chromosomes, where matches to *Fol* mobile chromosomes were concentrated. Future long-read sequencing allied to greater sampling of co-occurring taxa would confirm or refute this possibility.

Our results also provide the first insights into the potential genetic basis for pathogenicity in coffee wilt disease. Of 65 putative effector genes we identified, roughly one third were shared by all *F. xylarioides* isolates. These are expected to include genes that contribute to the common pathogenicity traits of these fungi, i.e. those effectors which migrate to and induce vascular wilting first in the main infected stem and latterly in the systemic infection of the coffee tree as visualised as as premature ripening of coffee berries (Flood 2009). The main differences that we detected between host-specific populations were in gene content and expansion of gene families. For example, while *Fusarium* is broadly enriched for lytic enzymes involved in carbohydrate metabolism (Reis et al. 2005, Soanes et al. 2008), and vascular wilt inducers for certain gene families (as also reported by Klosterman et al. (2011)), we found differences between the arabica and robusta populations, particularly in metabolism of plant cell wall components. Specifically, the host-specific populations had different enzymes although they share similar functions, for example the arabica strains share five CAZyme copies involved in the hydrolysis of chitin, pectin and rhamnose, whereas the robusta strains share twelve CAZymes with the *Coffea* strains that are involved in the breakdown of glucose, xylan and mannose (none are found just in the robusta strains). Whilst we cannot precisely infer the phenotypic effects of these changes, expanded cell-wall degrading enzymes imply variation in the capacity or mode of *F. xylarioides* to breakdown plant cell walls. Cell-wall breakdown releases pectins into the xylem vessels, which could unintentionally act as a barrier to further pathogen growth. Pectin-degrading enzymes may therefore be additionally important for breaking down pectin barriers for the pathogen to spread leading to the external symptoms of wilting as infection progresses (Clérivet et al. 2000). The ability to utilise plant carbon suggests the fungus is able to live within host coffee plants for a long time before the disease state becomes evident (Klosterman et al. 2011), something which has been reported from the field for *F. xylarioides* (Flood 2009). A key benefit of the availability of living samples is that we can now begin to test the importance of these differences with functional assays.

Finally, our results highlight the multigenic nature of how fungi respond to changing crops and host specialisation. The complexity of the process whereby new effectors emerge means that many pathogenicity genes have functional redundancy, whereby disruption of single genes does not affect virulence (Reignault et al. 2008). Therefore, it is expected that no single gene confers host specificity, but a diverse set of pathogenicity genes is required. In turn, the emergence of newly pathogenic or host-specific strains involves shifts in profiles of multiple genes. This can occur by transfer of whole pathogenicity regions or from the emergence of distinct lineages that diverged over longer periods of time, but later independently emerge as significant crop pathogens, as observed here for the arabica versus robusta populations. Our finding of the transfer of pathogenicity factors between more distantly related taxa raises a particular concern for the currently widespread practice of intercropping plant species which share closely related plant pathogens, which might increase the probability of new virulent pathogens emerging. These findings prove the value of culture collections in providing historic and optimally preserved strains which help us to understand and model the evolution and spread of disease.

## 5 Materials and Methods

### 5.1 Strain history

We sequenced the genomes of six strains of *F. xylarioides* (table 1). Two strains derive from the 1950s and the pre-1970s outbreak in CAR and Cote D’Ivoire respectively: *Coffea659* (IMI 127659/ DSMZ 62457) and *Coffea674* (IMI 392674/ CBS 258.52). We call these *Coffea* strains because of their ability to infect multiple *Coffea* species including robusta. *Coffea674* is the ex-type and in common with most pre-1970s strains infects robusta and other *Coffea* species but not arabica (Lepoint et al. 2005). *Coffea659* is one of the few strains able to also infect arabica in trials and therefore is a true host generalist (Girma 2004, Flood 2009). The remaining strains are grouped by their molecular markers (Buddie et al. 2015) and comprise two arabica host-specific strains (arabica563, IMI 389563 and arabica908, IMI 375908); and two robusta strains (robusta254, IMI 392254 and robusta277, IMI 392277), all collected five years apart between 1990-2000. At around the same time that CWD re-emerged on robusta coffee, it was also reported in Ethiopia on “arabica” coffee (*C. arabica*) (Stewart 1957, Lejeune 1958) and *F. xylarioides* was confirmed as the causal agent (Kranz & Mogk 1973, Van Der Graaff & Pieters 1978). By the 1990s, CWD was causing widespread destruction of arabica coffee in Ethiopia, and robusta coffee in northeast DRC, Uganda and northern Tanzania.

### 5.2 Strain information

Six *F. xylarioides* strains were selected for Illumina MiSeq sequencing (table 1). All strains were originally collected over a 52 year period from CWD-infected trees and stored in the CABI culture collection (CABI, Egham, Surrey, TW20 9TY). Strains were grown on synthetic low nutrient agar (SNA) at 25°C, and potato sucrose agar (PSA) with UV irradiation in order to confirm morphological identification following (Buddie et al. 2015). Following Cubero et al. (1999), strains were grown in GYM broth and genomic DNA was extracted from <20mg of washed mycelium, which was frozen in liquid nitrogen before extraction folllowing the DNeasy Plant Mini Kit (Qiagen, Hilden, Germany) standard protocol.

### 5.3 Genome sequencing and assembly

For each strain, a single library was prepared with the Illumina TruSeq PCR-free kit and sequenced with the Illumina MiSeq v3 600 cycle kit with 2 x 300 bp paired end sequencing and 350bp insert size at the Department of Biochemistry (University of Cambridge, UK). Low quality bases and adaptors were identified and trimmed (quality score <20) using TrimGalore 0.6.0 by Cutadapt (https://github.com/FelixKrueger/TrimGalore, Martin (2011)). Overlapping pairedend reads were merged using FLASH 1.2.11 (**?**) before being assembled using MEGAHIT 1.2.8 (**?**) with a final k-mer length of 189. Assembly metrics were computed using QUAST 5.0.2 (**?**) (table 1). The quality of the genomes in terms of the presence of 303 core eukaryotic genes and 290 core fungal genes was assessed using BUSCO v3.0.2 (Simão et al. 2015). All raw sequence data has been deposited in the relevant International Nucleotide Sequence Database Collaboration (INSDC) databases under the Study IDs ******* (see table S* for Run accessions). For comparison, we selected eleven genomes of related species (table S1) selected for being: closely related and wilt-inducers (*F. udum, F. oxysporum f. sp. lycopersici* and *f. sp. cubense*); closely related which occupy the same coffee-plant niche (*F. verticillioides, F. solani* and *F. oxysporum*); closely related (*F. fujikuroi, F. mangiferae, F. proliferatum*); and wilt-inducers (*V. dahliae* and *V. alboatrum*). Chromosomal and lineage-specific regions in each *F. xylarioides* assembly were identified by whole-genome alignment to the chromosomal level assembly of *F. verticillioides* (GenBank accession GCA 003316975.2, Ma et al. 2010). The *Coffea659* assembly (chosen for having the larger genome of the two *Coffea* strains) was aligned against *F. verticillioides* using RaGOO v1.1 with the ‘-C’ parameter (Alonge et al. 2019), with the remaining *F. xylarioides* strains aligned in-turn against the assembled RaGOO *Coffea659*. Scaffolds which did not match to *F. verticillioides* but are present in each *F. xylarioides* strain and the *F. udum* genome (GCA 002194535.1, Srivastava 2018) were interpreted as *Fusarium xylarioides* and udum-specific (FXU). Scaffolds which are present only in *F. xylarioides* were interpreted as *F. xylarioides*-specific, while scaffolds which did not match to the RaGOO *Coffea659* and *F. verticillioides* assembly nor *F. udum* were concluded to be lineage-specific (LS). Here, the LS scaffolds relate to the host-specific populations.

### 5.4 Gene prediction and orthologous clustering

Protein-encoding genes were predicted using the BRAKER v2.1.2 pipeline (Stanke et al. 2006, 2008, Li et al. 2009, Barnett et al. 2011, Hoff et al. 2016, 2019), using RNA-seq data from the closely related *F. verticillioides* to guide gene models. RNA-seq reads (SRR10097610) were mapped to each genome using STAR v2.7.3a (**?**) and the resultant BAM file was input to BRAKER with default settings. BUSCO analysis was performed to assess quality of predicted proteomes, and the GC content of coding sequences was calculated. This was used to calculate GC-equilibrated regions in 20 kb windows, where GC-rich blocks have a GC content within 1% of that of the coding sequences, as well as AT-rich blocks, where the GC content is ¡= 0.7 of the coding sequences content (Rouxel et al. 2011). A Kruskal-Wallis chi-squared test confirmed association between genes and the GC-rich blocks, with a Wilcoxon rank test (Bonferroni adjustment) to confirm significance between pairs. Three programs were used to infer functional classification: Interproscan v5.35-74.0 (Jones et al. 2014); Superfocus (**?**) which classifies the proteins by their SEED categories (Aziz et al. 2012); and NCBI BLAST (https://blast.ncbi.nlm.nih.gov/Blast.cgi) using megaBLAST searches and the nr database. All proteins lacking significant hits were annotated as hypothetical proteins. OrthoFinder v2.3.8 (Emms & Kelly 2017, 2019) was used to determine orthologous groups amongst the six *F. xylarioides* strains and eleven related species (table S1). Orthofinder also reconstructed a species tree from gene trees of the entire set of orthologs reconstructed by FastML (figure 4). Concordance of the set of individual gene trees for single ortholog groups was summarised using Phy-Parts (https://bitbucket.org/blackrim/phyparts/src/master/), while concordance of the ortholog groups was summarised using venn diagrams drawn with the Venn package in RStudio. Correlations between strains were tested for an excess of sharing using the “SuperExactTest” package in RStudio ((Wang et al. 2015).

### 5.5 Transposable elements and repeat annotation

Repeats and transposable elements (TEs) were identified from the assembled nucleotides using the RepeatModeler and RepeatMasker pipelines (**??**). The distribution of TEs across the genome was represented as the fraction of base pairs within 20kb windows assigned to TEs. In the same way as for genes, the association between TEs and AT-rich areas was tested using a Kruskal-Wallis chi-squared and a Wilcoxon rank test (bonferroni adjustment). The separate distribution of Class 1 RNA retrotransposons and Class 2 DNA transposons (Daboussi & Capy 2003) was also calculated. The presence of repeat-induced point mutations (RIP) were confirmed using The RIPper bioinformatics tool (Van Wyk et al. 2019), with default settings applied to identify Large RIP Affected Regions (LRARs), and the RIPCAL (Hane & Oliver 2008) RIP-index scan with default threshold settings applied and a scanning subsequence length of 10kb for the core chromosome scaffolds and 300bp for the FXU and LS scaffolds to reflect the average sequence length across the different scaffold groups.

### 5.6 Searching for putative effector genes involved in wilt disease

We adopted a four-pronged approach to search for putative effector genes involved in the host-specificity between the arabica and robusta populations.

Pre-characterised effectors. Putative effector sequences described in closely related *Fusarium* species were downloaded from Genbank (table S3). Using BLAST, the sequences were searched for against the *F. xylarioides* predicted genes and scaffolds. Proteins were judged to be present if a blast match of >70% and 1.00E-50 was obtained and the sequence retrieved from the genome scaffold, checking to include the full protein region if the blast match only returned a partial match.

Small, cysteine-rich putative effectors. Many fungal effector proteins are small (¡400 amino acids), cysteine-rich (¿4 cysteine residues) and secreted (Van Esse et al. 2008, de Jonge et al. 2011). Putative effectors were therefore searched for within each genome based on size, cysteine-richness, secretion signal, putative function and genome locus. First, annotated genes were sorted according to size and number of cysteine residues using SeqKit (Shen et al. 2016), to select those with ¡400 amino acids and ¿4 cysteine residues. Next, the presence of a signal peptide and a signal peptide cleavage site on those proteins were predicted using TargetP 1.01 (Emanuelsson et al. 2000, Nielsen et al. 1997) with an RC score cut-off 1-3 to increase specificity. Finally, subcellular localizations for these proteins were predicted using WoLF PSORT (https://wolfpsort.hgc.jp/) to identify secreted extra-cellular proteins. The orthologous gene sets for those genes were identified from the Orthofinder results.

CAZyme effectors. CAZymes are carbohydrate-active enzymes thought to be important in the infection pathway. Carbohydrates in plant cell walls provide the main source of carbon for fungal pathogens (Hervé et al. 2010) and consist of cellulose microfibrils embedded in a matrix of hemicelluloses, pectin polysaccharides and glycoproteins (Carpita & Gibeaut 1993). The breakdown of cell wall polymers requires a range of enzymes, including glycoside hydrolases (GHs), pectate and polysaccharide lyases and carbohydrate esterases (CEs) (Mace et al. 1981). CAZymes that target cell walls also contain carbohydrate-binding modules (CBMs) (Boraston et al. 2004), which bind to cell wall polymers and increase the enzymes’ catalytic efficiency by improving contact (Hervé et al. 2010, **?**). CBMs show specificity for particular polymers (McCartney et al. 2006). Broadly, *Fusarium* is enriched for CAZymes (table S4). Therefore, to identify putative CAZyme-encoding effector genes, and the CBM and their associated carbohydrate active modules differentially expressed across the wilt-inducing and non-wilt inducing fungal strains we used the CAZy database (www.cazy.org, Lombard et al. (2013)) to identify CAZyme-encoding orthologous groups. We looked for those present in only the wilt-inducing strains using BLAST (e-vale 1.00e-50), additionally allowing for those which were present in just one non-wilt inducing strain.

Transposon-adjacent. In particular, class 2 MITES (Miniature Inverted-repeat Transposable Elements) have been shown to associate with effector proteins (Schmidt et al. 2013). The abundance of miniature and full impala transposons, as well as recently described transposons across the genomes was tested using the accessions detailed in table S7, with a BLAST (score >70% and 1.00e-50) confirming their presence.

For each set of putative effectors, nucleotide sequences were aligned using MAFFT and maximum likelihood trees reconstructed using PhyML 3.0 (Guindon et al. 2009) with a Generalised Time-Reversible model with invariant sites and gamma distributed variation in substitution rates across 4 rate classes of sites, implemented in Geneious v9.1. We also located each putative effector on the genome scaffolds in each species mapped against the chromosomal level *F. verticillioides* assembly. The site test of positive selection in PAML v4.8 (Yang 2007) was used to test for positive selection among codons in the DNA sequences in each phylogenetic tree by estimating the ratio between synonymous and nonsynonymous substitutions (*ω*). PAML detects positive selection when is significantly greater than 1 for a subset of codons. We compared a null model of “NearlyNeutral” selection that includes a class of codons under purifying selection and a class that evolve neutrally, against an alternative model of “PositiveSelection” that additionally includes a class with >1. Log-likehood ratios were calculated and compared with a chi-squared test with one degree of freedom to confirm significance. A Bayes Empirical Bayes (BEB) approach was used to identify specific amino acid sites under positive selection within gene sequences which were confirmed to be under positive selection (Yang et al. 2005).

## 6 Acknowledgments

This research was supported by the Natural Environment Research Council [grant number NE/L002515/1] (LP) and NE/S010866/1 (TB and RN). With thanks to CGR and NERC Metagenomics Workshop and with special thanks to H. Stewart and A. Buddie (CABI) for invaluable lab support and advice, and C. Wilson, P. Spanu, B. Murphy and T. LLewellyn for comments on the manuscript.

