## Supplementary material for "Historical genomics reveals the evolutionary mechanisms behind multiple outbreaks of the host-specific coffee wilt pathogen *Fusarium xylarioides*"

### Supplementary figures

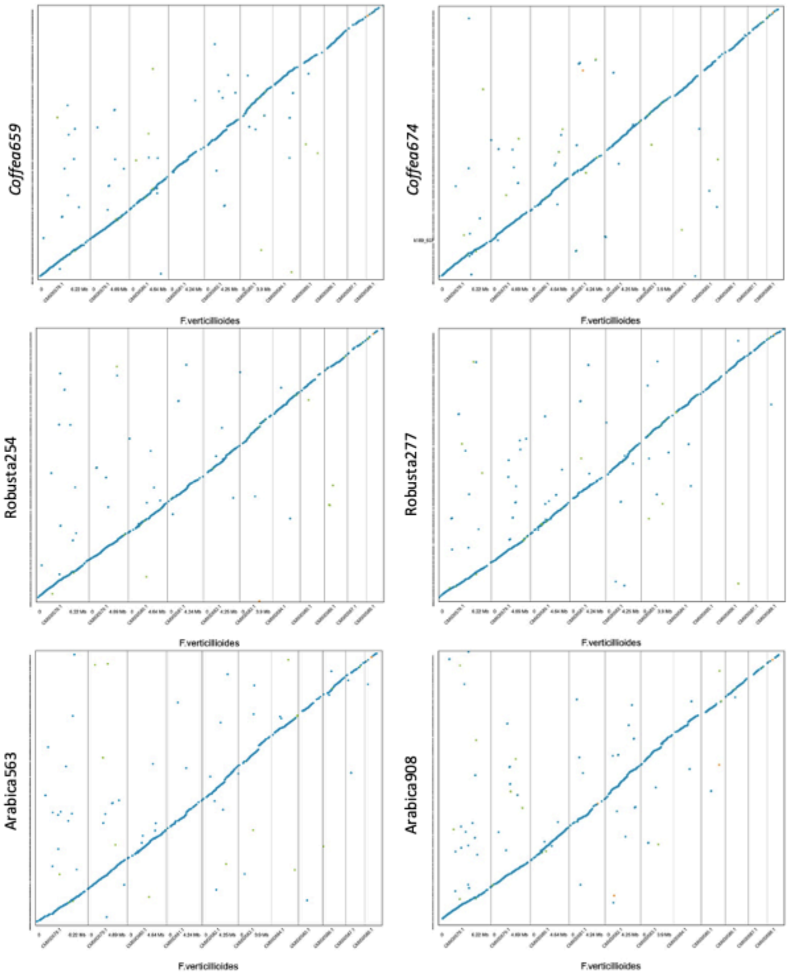


Supplementary figure 1. Representative whole-genome alignments of *F. xylarioides* strains against the 11 *F. verticillioides* core chromosomes. Each dot represents chromosomal correspondence. Genomes were aligned using Mummer 4.0.0, with outputs processed using DotPrep.py before visualizing using Dot in DNANexus. Blue indicates forward alignments, green indicates reverse alignments, orange indicates repetitive alignments.

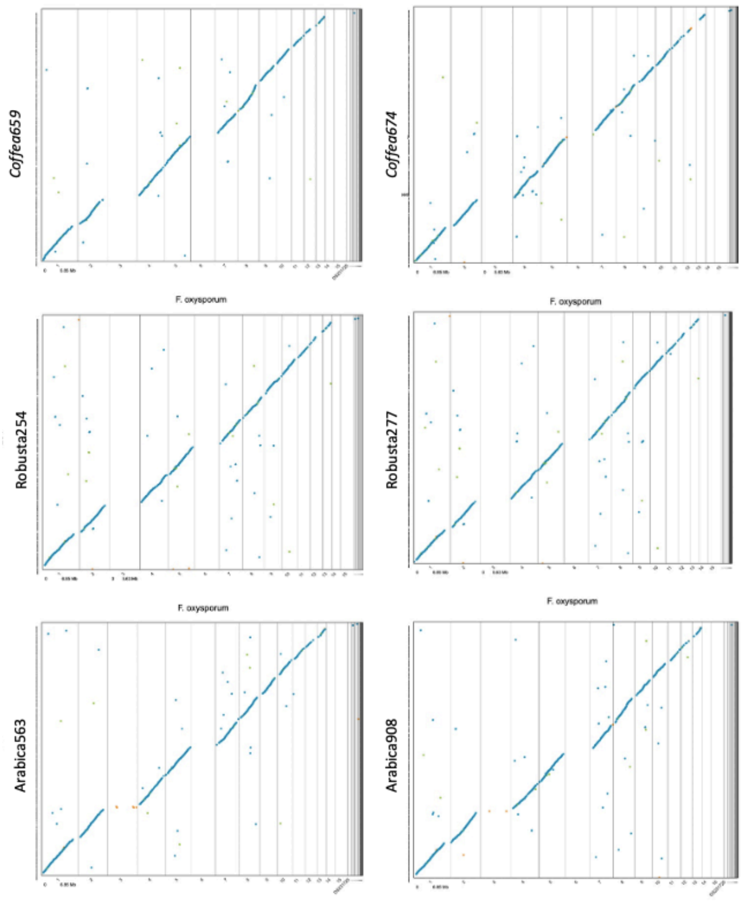


Supplementary figure 2. Representative whole-genome alignments of *F. xylarioides* strains against the 15 *F. oxysporum f. sp. lycopersici (Fol)* chromosomes, including 11 core chromosomes shared with *F. verticillioides* and 4 mobile chromosomes. Each dot represents chromosomal correspondence, with absences representing the absent *Fol* chromosomes. Genomes were aligned using Mummer 4.0.0, with outputs processed using DotPrep.py before visualizing using Dot in DNANexus. Blue indicates forward alignments, green indicates reverse alignments, orange indicates repetitive alignments.

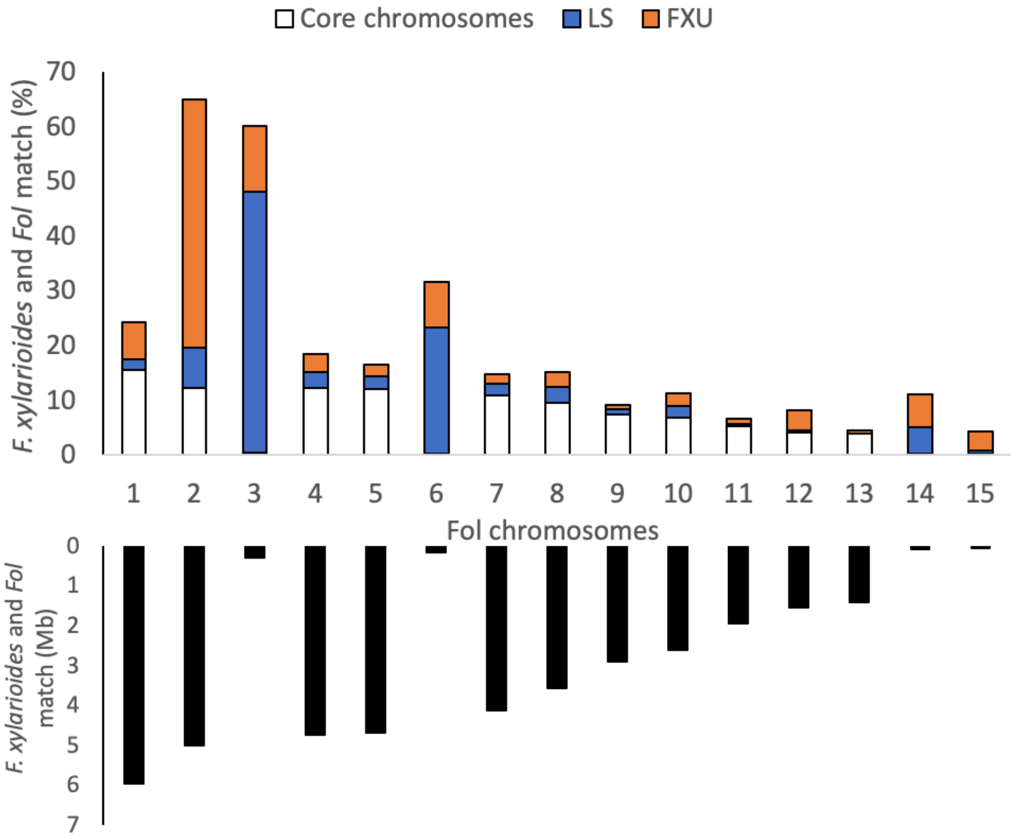


Supplementary figure 3. The association between *F. xylarioides* and *F. oxysporum f. sp. lycopersici (Fol)* from reference-guided scaffolding. A The proportion of *F. xylarioides* scaffolds which match a *Fol* chromosome by scaffold group: core chromosome; lineage-specific (LS); and *F. xylarioides*- and *-udum* specific (FXU). B The total match for each *Fol* chromosome with *F. xylarioides.*

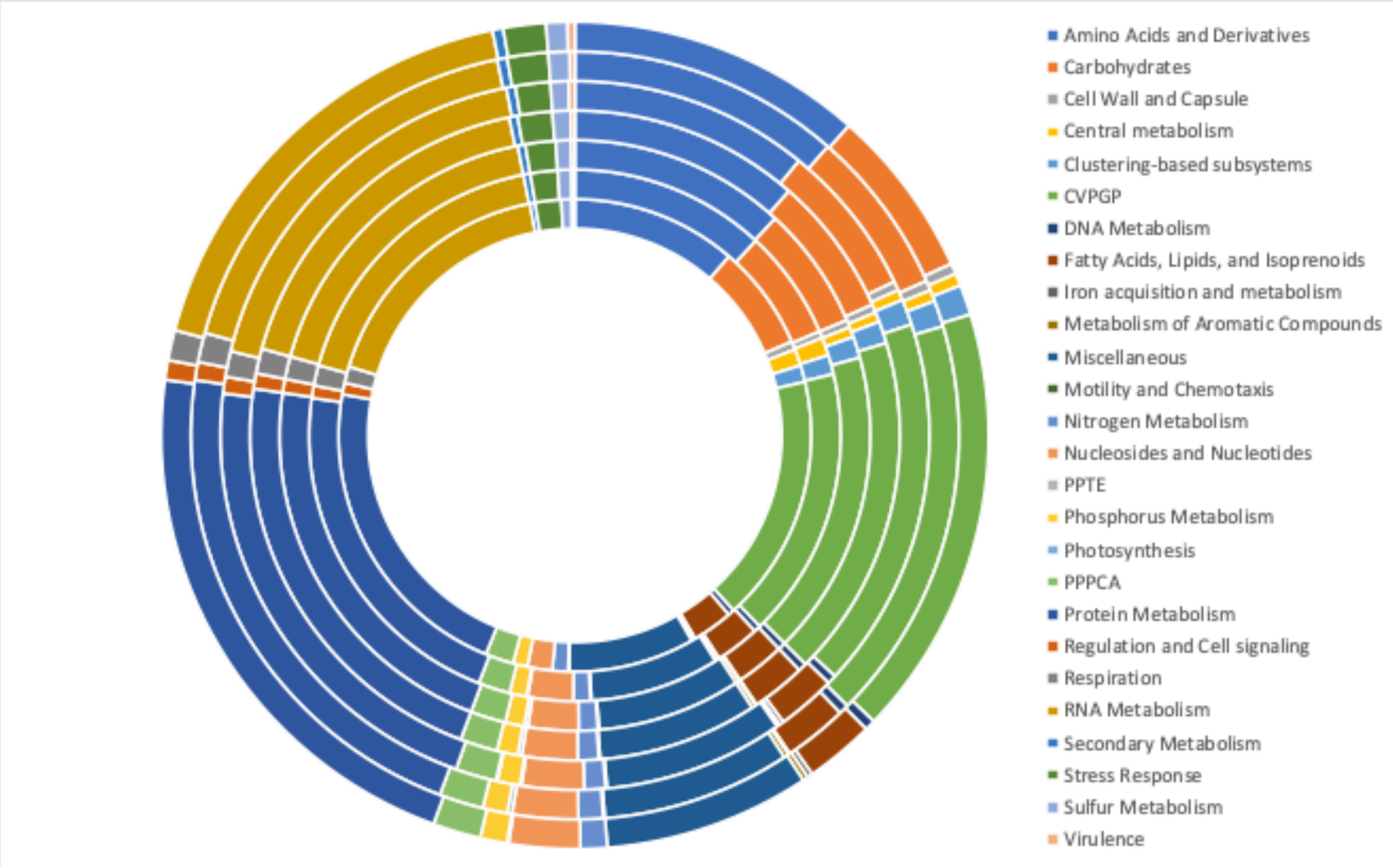


Supplementary figure 4. Functional diversity of *F. xylarioides* and *F. udum* (from inner circle outwards: F. *udum*; robusta254; robusta277; arabica563; arabica908; *Coffea659; Coffea674*). Classification was based on the number of hits to each Level 1 category in SUPER-FOCUS (Silva et al. 2017). Abbreviations: CVPGP = Cofactors, Vitamins, Prosthetic Groups, Pigments; PPTE = Phages, Prophages, Transposable elements; PPPCA = Predictions based on plant-prokaryote comparative analysis

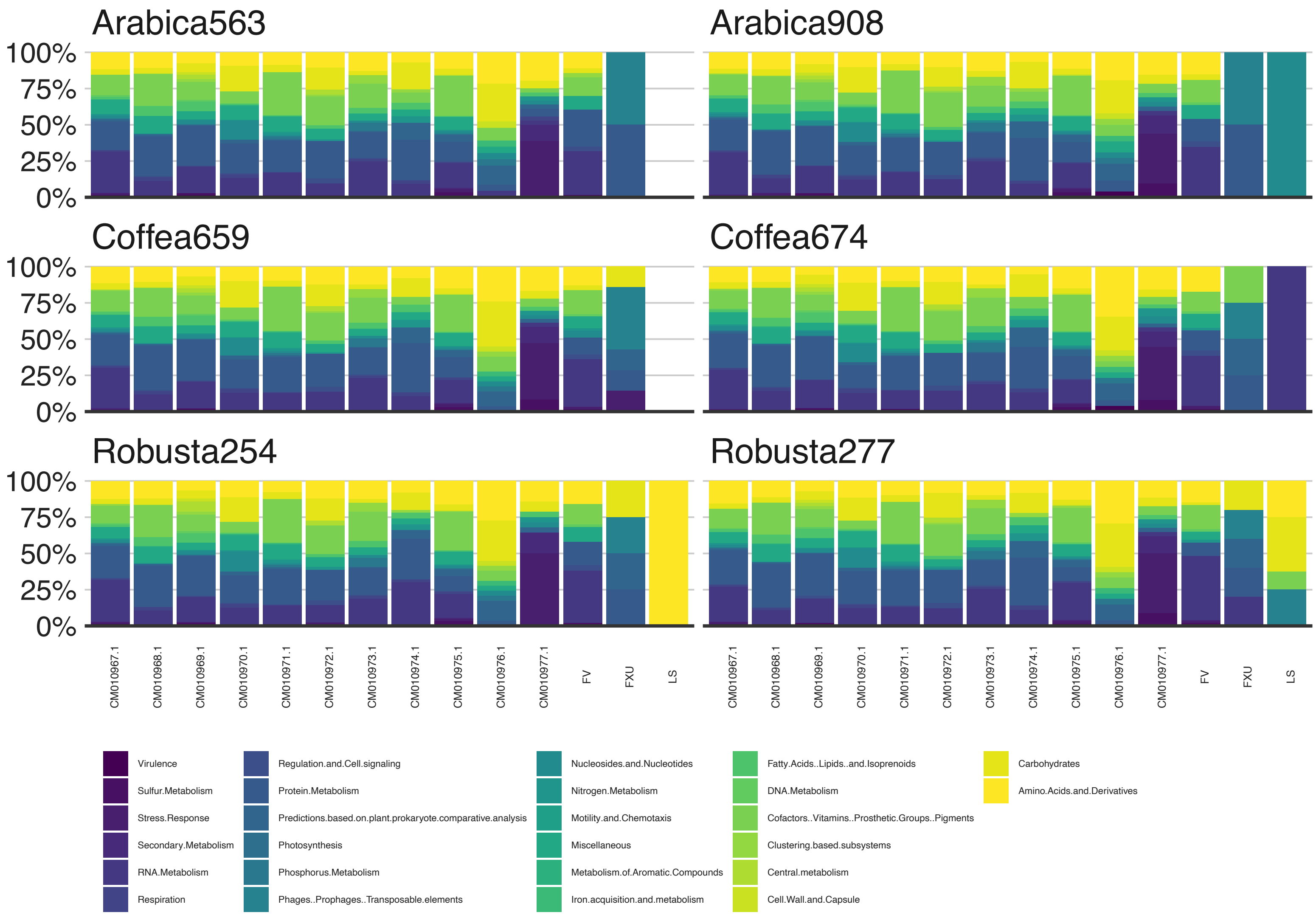


Supplementary figure 5. Functional diversity of *F. xylarioides* across its core chromosomes and by scaffold group: lineage-specific (LS); *F. xylarioides*- and -udum-specific. Classification was based on the number of hits to each Level 1 category in SUPER-FOCUS (Silva et al. 2017)

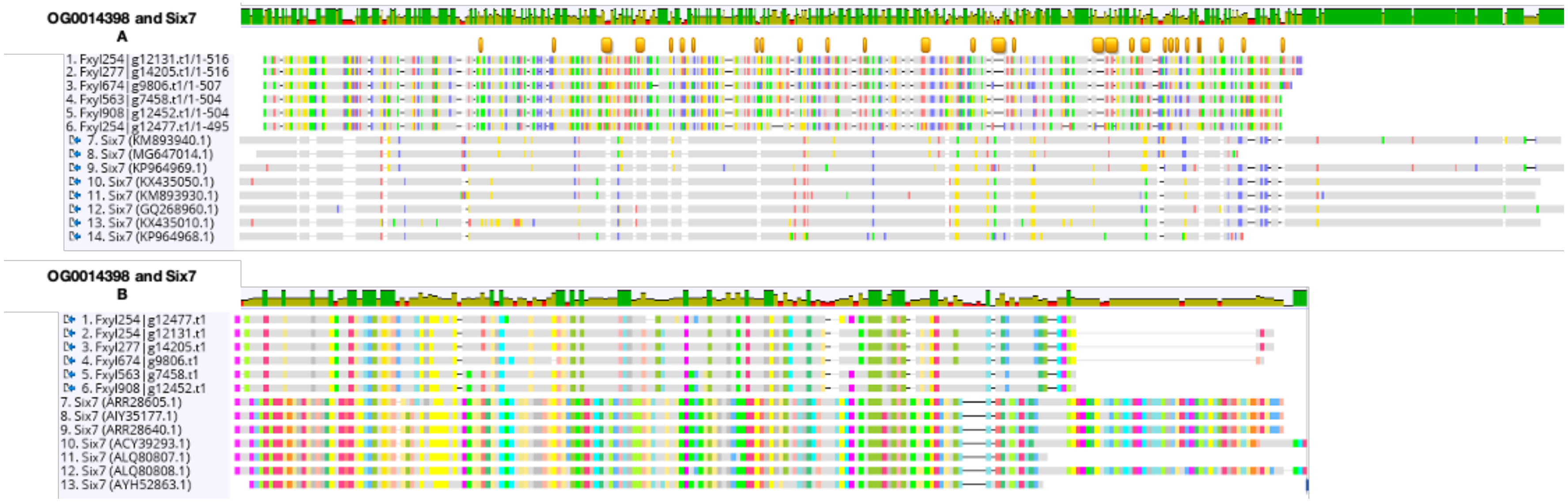


Supplementary figure 6. A A nucleotide alignment between the orthogroup OG0014398 and seven *six7 F. oxysporum* nucletoide sequences, and B A protein alignment between the orthogroup OG0014398 and seven Six7 *F. oxysporum* amino acid sequences. Sequences aligned with MAFFT and drawn in Geneious 9.1.

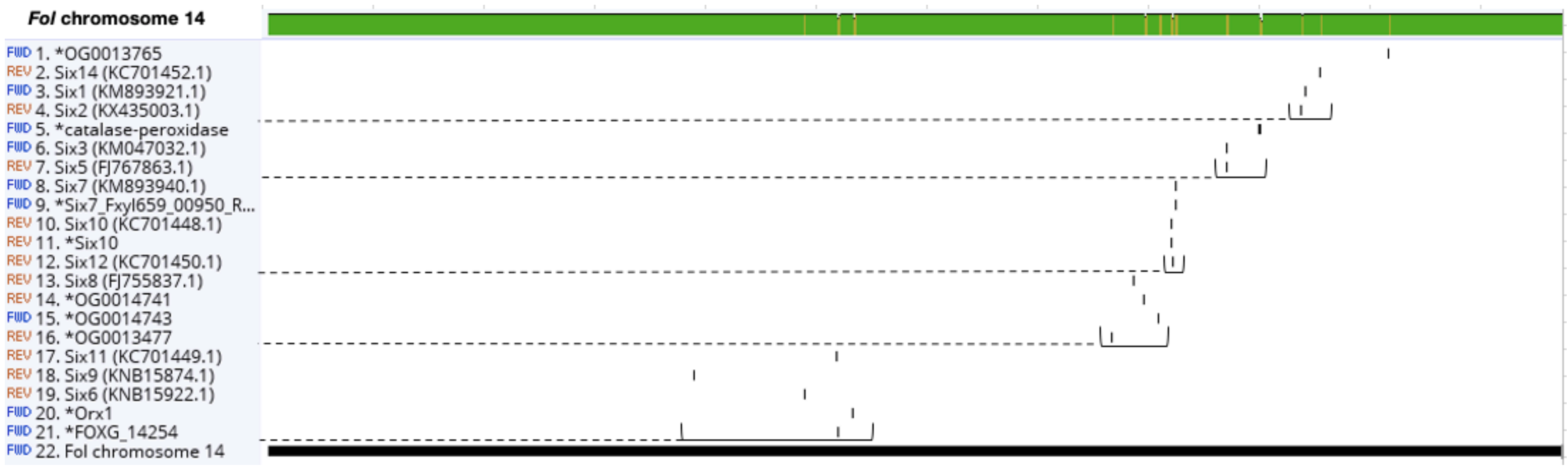


Supplementary figure 7. *F. oxysporum f. sp. lycopersici (Fol)* chromosome 14 aligned with all known SIX effectors and the nine effectors described in this study which match regions of chromosome 14 (pre-fixed with a *). Five SIX chromosomal mini-clusters (described by Schmidt et al. (2013) are marked on the genome plot. Eight effectors all reside close to four of these clusters: *FOXG*_14254 and *orx1* are 1.8 kb and 24 kb from *six11* (which is clustered with *six6* and five genes (including *orx1* in *Fol*) and a transcription factor); OG0013477, OG0014741 and OG0014743 are 30 kb, 12.5 kb and 33 kb respectively from *six8* (which resides in a solo block with class two transposons and a gene encoding an unknown protein); *six7* (from 659_00950) shares a locus with *Fol six7* and its cluster with *six10* and *six12*. The orthogroups OG0014741 and OG0014743 are both also <40 kb from this *six10, six12, six7* minicluster. Finally, *catalase-peroxidase* is 45 kb from the *six3, six5* minicluster. OG0013765 is 95 kb from the *six1, six2, six14* minicluster. Sequences aligned with MAFFT and drawn in Geneious 9.1.

| | Robusta | | Coffea | | Arabica | | | Transposon (bp from promoter) | Subcellular location | Closest species match (BLAST) | F. oxysporum percent identity (%) |
|---|---|---|---|---|---|---|---|---|---|---|---|
| Protein | Fx254 | Fx277 | Fx659 | Fx674 | Fx563 | Fx908 | Fud | | | | |
| fow1 | | | | | | | | | cyto | F. fujikuroi | |
| pelD | ^ | ^ | ^ | ^ | ^ | ^ | | | extr | F. verticillioides | |
| fmk1 | | | | | | | | | nucl | F. oxysporum f. sp. lycopersici | |
| **sge1** | | | | | | | | | nucl | F. acutatum | |
| snf1 | | | | | | | | | nucl | F. verticillioides | |
| **pep1** | | | | | | | | | | | |
| chlo_vacu | ^ | ^ | ^ | ^ | ^ | ^ | | | pero | F. oxysporum | |
| rho1.1 | | | | | | | | | cyto | F. oxysporum f. sp. lycopersici | |
| rho1.2 | | | | | | | | | nucl | F. pseudograminearum | |
| pelA | ^ | ^ | ^ | ^ | ^ | ^ | | | extr | F. oxysporum f. sp. lycopersici | |
| FOXG_14254 | | | ^ | | ^ | ^ | | | extr | F. anthophilum | |
| Orx1 | ^ | ^ | ^ | ^ | ^ | ^ | | | extr | F. acutatum | |
| catalase-peroxidase | ^ | ^ | ^ | ^ | ^ | ^ | | | extr | F. fujikuroi | |
| nep1 | | | | | | | | | mito | F. fujikuroi | |
| glucosyltransferase | | | | | | | | | nucl | F. fujikuroi | |
| pda1 | " | " | | | | | | | nucl | F. oxysporum f. sp. pisi | 89 |
| Six7 | | | ^ | | ^ | ^ | | | | | |
| **Six10** | | | | | | | | | nucl | F. oxysporum | |
| cytoskeletal | | | | | | | | | cysk | F. oxysporum f. sp. pisi | |
| OG0013792 | | | | | | | | | plas | F. oxysporum | 40 |
| OG0013889 | ^ | ^ | ^ | ^ | ^ | ^ | | | extr | F. oxysporum | 85 |
| OG0013871 | ^" | ^" | ^" | ^" | ^" | ^" | | | extr | F. oxysporum | 90 |
| **OG0013477** | ~^" | ~^" | ~^" | ~^" | ~^" | ~^" | | | extr | F. oxysporum f. sp. pisi | 94 |
| OG0013861 | * | * | * | * | * | * | * | | | F. poae | |
| OG0013645 | | | | | | | | | | F. proliferatum | |
| OG0013763 | ^ | ^ | ^ | ^ | ^ | ^ | | | cyto_extr | Neonectria ditissima | |
| OG0013877 | ^ | ^ | ^ | ^ | ^ | ^ | | | extr | F. langsethiae | |
| OG0013738 | ~^ | ~^ | | ~^ | ~^ | ~^ | | | extr | F. mangiferae | |
| OG0014864 | | | | | | | | | plas | F agapanthi | |
| **OG0014398/Six7** | * ~^ | * ~^ | | * ~^ | * ~^ | * ~^ | | 587 | extr | F. oxysporum | 63 |
| OG0014238 | ~" | " | | ~" | " | ~" | | | cyto | F. oxysporum f. sp. vasinfectum | 92 |
| OG0014828 | | *^ | | *^ | * | * | * | | | F. oxysporum | 77 |
| OG0014165 | *^ | *^ | *^ | *^ | | | | | extr | F. oxysporum f. sp. radici-cucunerinum | |
| OG0015453 | ^ | ^ | ^ | ^ | | | | | | F. beomiforme | |
| OG0014836 | ^ | ^ | | ^ | | | | | | F. oxysporum f. sp. radici-cucunerinum | 77 |
| OG0015372 | *^ | *^ | | *^ | | | | | extr | F. oxysporum | |
| OG0016234 | ^ | ^ | ^ | | | | | | extr | - | |
| OG0013787 | | | ^ | ^ | ^ | ^ | | | | F. acutatum | |
| OG0016323 | | | | | | | | | plas | F. napiforme | |
| OG0014797 | *^ | *^ | | | *^ | *^ | | | extr | F. anthophilum | |
| OG0014367 | | | | | | | | | | F. agapanthi | |
| OG0013912 | ^ | ^ | ^ | | ^ | ^ | | | extr | F. nygamai | |
| OG0008649 | " | " | " | " | " | " | | | cyto | F. oxysporum | 94 |
| OG0014891 | | | | | | | | | cysk | F. oxysporum | 82 |
| OG0016261 | | | | | | | | | pero | F. oxysporum f. sp. cubense | 78 |
| OG0016232 | | | | | | | | | | Trichoderma harzianum | |
| **OG0016241** | | | | | | | | | mito | F. oxysporum | 84 |
| OG0016247 | ~" | " | | ~" | | | | | | F. oxysporum | 93 |
| OG0014811 | ^ | ^ | ^ | ^ | | | | | extr | F. oxysporum f. sp. raphani | 83 |
| OG0015465 | ~ | ~ | ~ | ~ | | | | | mito | F. nygamai | |
| **OG0014741** | *" | *" | *" | *" | | | | | | F. oxysporum f. sp. raphani | 99 |
| **OG0014743** | " | " | " | " | | | | 389 | plas | F. oxysporum f. sp. raphani | 99 |
| OG0016212 | | | | | ~^" | ~^" | | | extr | F. oxysporum f. sp. cubense | 98 |
| OG0018569 | | | | | ~" | ~" | | | cyto_nucl | F. oxysporum | 92 |
| OG0018533 | | | | | | | | | cyto | F. fujikuroi | |
| OG0018547 | | | | | | | | | cyto_mito | F. fujiuroi | |
| OG0018531 | | | | | | | | | mito | F. oxysporum | 79 |
| OG0014180 | | | | | *^" | *^" | | | extr | F. oxysporum | 90 |
| **OG0014392** | *~ | *~ | *~ | *~ | *~^ | *~^ | | 0 | | F. oxysporum f. sp. pisi | 48 |
| **OG0013478** | " | " | " | " | | | | 448 | extr | F. oxysporum f. sp. raphani | 98 |
| **OG0009441** | | | | | | | | 242 | mito | F. fujikuroi | |
| **OG0000973** | | | | | | | | 1035 | plas | F. oxysporum f. sp. vasinfectum | |
| **OG0013765** | | | | | | | | 0 | nucl | F. oxysporum f. sp. cepae | |
| **OG0015458** | " | " | " | " | | | | 0 | nucl | F. oxysporum | 99 |
| **OG0000409** | | | | ^ | ^ | ^ | | | extr | F. verticillioides | |
| **OG0014179** | | | | | " | " | | 1291 | | F. oxysporum f. sp. radici-lycopersici | 97 |

Supplementary figure 8. Putative effectors' characteristics and presence or absence across *F. xylarioides* strain and *F. udum*. The four effector classes are shown in: yellow for predefined effectors; purple for small and cysteine-rich secreted effectors; blue for carbohydrate-active enzymes; and red for transposon-adjacent effectors. The symbols highlight: the presence of transposons is represented by names in bold with its distance from the genes promoter described if less than 1500bp (if not, the transposon is over 1500bp away); genes under positive selection by an asterisk; genes in an AT-rich region by a tilde; genes with evidence of horizontal transfer from *F. oxysporum* are a darker shade; genes which are absent from more closely-related *Fusarium* species (namely *F. graminearum*, the Asian clade GFC species and F. *verticillioides* - *F. solani* and *F. udum* were excluded here because *F. solani* also infects coffee (Rutherford & Flood 2005) and thus could be a source of pathogenicity and *F. udum* is also a vascular wilt-inducer) and *F. oxysporum* is the closest match with a percent identity (%) >=90 are represented by a quotation mark; genes which share a locus across all strains are outlined in black; and closest species is shown for each protein with its percent identity (%), where a BLASTp hit was returned.

### Supplementary tables

Supplementary table 1. Published genomes used for comparison

| Genome | Accession number |
|---|---|
| *F. udum* | NIFK00000000.1 |
| *F. oxysporum f.sp. lycopersici* | AAXH00000000.1 |
| *F. oxysporum f.sp. cubense* | SRMI00000000.1 |
| *F. verticillioides* | AAIM00000000.2 |
| *F. fujikuroi* | GCA_900079805.1 |
| *F. mangiferae* | FCQH00000000.1 |
| *F. solani* | NGZQ00000000.1 |
| *Verticillium dahliae* | ABJE00000000.1 |
| *V. albo-atrum* | NMXJ00000000.1 |
| *F. graminearum* | ASM24013v3 |
| *F. proliferatum* | GCF_900067095.1 |

Supplementary table 2. Total interspersed repeats and transposon sequence lengths across *F. xylarioides, F. udum, Fol* and *F. verticillioides*. Transposons and repeats were identified and classified using the RepeatModeler and RepeatMasker pipelines (see methods).

| **Class** | **Family** | ***Coffea674*** | ***Coffea659*** | **Robusta254** | **Robusta277** | **Arabica563** | **Arabica908** | ***F. udum*** | ***Fol*** | ***F. verticillioides*** |
|---|---|---|---|---|---|---|---|---|---|---|
| **DNA** | hAT-Ac | 726,547 | 76,885 | 95,222 | 94,583 | 91,394 | 110,906 | 6,958 | | |
| | hAT-hobo | 35,501 | 56,812 | 7,056 | 7,056 | 1,137 | | | | |
| | MULE-MuDR | 25,895 | | | | 12,546 | 16,592 | 260,285 | 503,173 | |
| | PIF-Harbinger | 306,039 | 504,602 | 162,764 | 155,913 | 414,305 | 409,108 | 34,928 | | |
| | PiggyBac | 286,500 | 324,012 | 255,817 | 253,457 | 327,944 | 262,638 | | 229,376 | |
| | TcMar-Fot1 | 478,461 | 524,886 | 251,531 | 248,862 | 230,454 | 195,105 | 105,293 | 1,334,911 | 63,426 |
| | TcMar-Tc1 | 272,564 | 269,731 | 428,906 | 421,652 | 300,252 | 303,742 | 38,757 | 134,846 | |
| | Crypton-H | | | | | | | | 41,016 | |
| | Kolobok-H | | | | | | | | 40,957 | |
| | Unspecified | 80,792 | | 16,280 | 16,079 | 130,554 | 93,277 | | 616,319 | |
| | **Total** | **2,212,299** | **1,756,928** | **1,217,576** | **1,197,602** | **1,508,586** | **1,391,368** | **446,221** | **3,796,214** | **63,426** |
| **Low complexity** | Unspecified | 52,676 | 98,842 | 61,178 | 60,747 | 61,824 | 59,114 | 44,401 | 31,755 | 30,337 |
| | **Total** | **52,676** | **98,842** | **61,178** | **60,747** | **61,824** | **59,114** | **44,401** | **31,755** | **30,337** |
| **LTR** | Copia | 2,935,958 | 1,473,220 | 1,604,519 | 1,497,945 | 1,412,484 | 1,038,622 | 2,528,027 | 969,632 | |
| | Gypsy | 161,544 | 911,254 | 1,180,369 | 1,059,865 | 1,484,751 | 2,387,885 | 5,706,226 | 262,884 | 158,720 |
| | Unknown | | | | | | | 70,960 | | |
| | **Total** | **3,097,502** | **2,384,474** | **2,784,888** | **2,557,810** | **2,897,235** | **3,426,507** | **8,305,213** | **1,232,516** | **158,720** |
| **rRNA** | Unspecified | 26,911 | 274,761 | 63,534 | 63,347 | 128,175 | 121,767 | 29,680 | 257,079 | |
| | **Total** | **26,911** | **274,761** | **63,534** | **63,347** | **128,175** | **121,767** | **29,680** | **257,079** | |
| **Satellite** | centr | | | 3,427 | 2,093 | | | | | |
| | | | | **3,427** | **2,093** | | | | | |
| **Simple repeat** | Unspecified | 363,946 | 370,605 | 373,360 | 371,041 | 383,258 | 352,347 | 310,907 | 244,493 | 247,530 |
| | **Total** | **363,946** | **370,605** | **373,360** | **371,041** | **383,258** | **352,347** | **310,907** | **244,493** | **247,530** |
| **tRNA** | Unspecified | 1,850 | 5,161 | 2,131 | 2,131 | | 1,875 | 2,051 | 3,532 | 5,422 |
| | **Total** | **1,850** | **5,161** | **2,131** | **2,131** | | **1,875** | | | |
| **Unknown** | Helitron-2 | | | | | 12,827 | | | 21,326 | |
| | Unspecified | 7,561,578 | 9,984,881 | 11,642,532 | 11,040,009 | 13,634,728 | 12,468,428 | 3,457,228 | 3,809,318 | 518,665 |
| | **Total** | **7,561,578** | **9,984,881** | **11,642,532** | **11,040,009** | **13,647,555** | **12,468,428** | **3,457,228** | **3,809,318** | **518,665** |
| **Grand Total** | **Total** | **13,316,762** | **14,875,652** | **16,148,626** | **15,294,780** | **18,626,633** | **17,821,406** | **12,593,650** | **9,643,175** | **1,018,678** |

Supplementary table 3. Predefined effector protein genes analysed in this study. Genes marked with an * show predicted roles and locations only

| Name | Role | Query Accession | Query species | Length | Reference |
|---|---|---|---|---|---|
| FOXG_02706.2 | Glucosyltransferase* | KNA98333.1 | *Fol* | 1493 | (44) |
| FOXG_10732.2 | Cytoskeletal* | KNB10567.1 | *Fol* | 449 | (44) |
| FOXG_04660.2 | Chloroplast/ vacuole* | KNB01401.1 | *Fol* | 797 | (44) |
| Nep1 | Microbial elicitors of plant necrosis | AF036580.1 | *Foe** | 2617 | (67) |
| Fmk1 | MAP kinase | KC257048.1 | F. oxysporum | 603 | (19) |
| Fow1 | Mitochondrial carrier protein | KC134256.1 | F. oxysporum | 725 | (39) |
| Pda1 | Pisatin demethylase | KR855811.1 | F. oxysporum | 455 | (95) |
| PelA | Pectate lyase | MK918256.1 | *Fol* | 539 | (71) |
| PelD | Pectate lyase | KC294608.1 | F. proliferatum | 552 | (71) |
| Pep1 | Pea pathogenicity protein | EU436568.1 | Fusarium sp. | 216 | (33) |
| Rho1 | Rho GTP-ase activating protein | KC017411.1 | F. oxysporum | 665 | (56) |
| Sge1 | SIX (secreted in xylem) gene expression 1 | LC369105.1 | *For** | 565 | (59) |
| Snf1 | Protein kinase sucrose non-fermenting | KU048959.1 | F. commune | 625 | (66) |
| FOXG_14254 | Conserved secreted protein | KNB15932.1 | *Fol* | 1592 | (53) |
| Orx1 | In-planta secreted oxidoreductase enzyme | KNB15937.1 | *Fol* | 1860 | (53) |
| Catalase-peroxidase | Secreted enzyme | KNB19974.1 | *Fol* | 2385 | (53) |
| SIX10 | Secreted in xylem 10 | KNB20462.1 | *Fol* | 736 | (53) |

*Abbreviations for *F. oxysporum formae speciales* sister species: *Fol, F. oxysporum f. sp. lycopersici; Foe, F. oxysporum f. sp. erythroxyli; For, F. oxysporum f. sp. ricini*

Supplementary table 4. Enriched CAZyme gene familes across *Fusarium* wilt- (*F. xylarioides, F. udum, Fol)* and non-wilt inducing (*F. verticillioides, F. fujikuroi, F. graminearum*) genomes, compared with three other ascomycete fungi (*Trichoderma reesei, Aspergillus nigris and Magnaporthe grisea*)

| **Species** | **AA*** | **CBM*** | **CE*** | **GH*** | **GT*** | **PL*** |
|---|---|---|---|---|---|---|
| *Fol* | *165* | *330* | *70* | *746* | *396* | *35* |
| *F. xylarioides (Coffea*674*)* | *111* | *232* | *62* | *488* | *248* | *28* |
| *F. udum* | 119 | 228 | 63 | 503 | 257 | 30 |
| *F. verticillioides* | 130 | 265 | 66 | 597 | 336 | 28 |
| *F. fujikuroi* | 105 | 220 | 61 | 476 | 274 | 29 |
| *F. graminearum* | 92 | 191 | 52 | 394 | 241 | 27 |
| *Trichoderma reesei* | 57 | 127 | 25 | 304 | 196 | 8 |
| *Aspergillus nigris* | 94 | 151 | 55 | 434 | 321 | 10 |
| *Magnaporthe grisea* | 118 | 207 | 60 | 378 | 258 | 9 |

Supplementary table 5. Gene copy number for CAZyme-encoding orthologous groups shared across the vascular wilt-inducing *Fusarium* and *Verticillium* strains. Groups which also included one non-vascular wilt inducer were additionally included, and those which are also a putative effector are shaded the same colour as in figure 5. Where a species has a gene in the orthologous group which is not recognised as a CAZyme is represented with an asterisk.

| CAZyme | *F. graminearum* | *F. mangiferae* | *F. proliferatum* | *F. solani* | *F. verticillioides* | *F. udum* (Vascular wilt-inducers) | Robusta254 | Robusta277 | Coffea659 | Coffea674 | Arabica563 | Arabica908 | *Fol* | *Foc* | *V. albo-atrum* | *V. dahliae* | CAZy Genbank accession |
|---|---|---|---|---|---|---|---|---|---|---|---|---|---|---|---|---|---|
| CBM18 | | | | | OG0013846 | | OG0013846 | OG0013846 | OG0013846 | OG0013846 | OG0013846 | OG0013846 | | | | | |
| CBM38 | | | | OG0008649 | | OG0008649 | OG0008649<br>OG0008649 | OG0008649<br>OG0008649 | OG0008649 | OG0008649 | OG0008649 | OG0008649 | OG0008649<br>OG0008649<br>OG0008649 | OG0008649 | OG0008649 | OG0008649 | |
| | | | | | | | | | | | OG0018547 | OG0018547 | | | | | *F. fujikuroi* |
| CBM50 | | | | | | | OG0013477 | OG0013477 | OG0013477 | OG0013477 | OG0013477 | OG0013477 | OG0013477 | | | | *Fol* |
| | | | | | OG0013846 | | | | | | OG0013846 | OG0013846 | | | | | *Cordyceps militaris* |
| CBM66 | | | | | | OG0012516 | OG0012516 | OG0012516 | OG0012516 | OG0012516 | OG0012516 | OG0012516 | OG0012516<br>OG0012516<br>OG0012516 | OG0012516 | | | *Bacillus* |
| GH35 | | OG0012085 | | | | OG0012085 | OG0012085 | OG0012085 | OG0012085 | OG0012085 | OG0012085 | OG0012085 | OG0012085 | OG0012085<br>OG0012085 | | | |
| | | | | | | | | | | | OG0018531 | OG0018531 | | | | | |
| GH16 | | | | | | | OG0013799 | OG0013799 | OG0013799 | | OG0013799 | OG0013799 | OG0013799 | OG0013799 | | | |
| GH28 | | | | | | | OG0014847 | OG0014847 | OG0014847 | | OG0014847 | OG0014847 | | | | | *F. oxysporum* |
| GH29 | | | | | | OG0013912 | OG0013912 | OG0013912 | OG0013912 | OG0013912 | OG0013912 | OG0013912 | | | | | |
| GH31 | | | | | | | OG0016261 | OG0016261 | OG0016261 | | | | | | | | |
| | | | | | | | | | | | OG0018569 | OG0018569 | | | | | *Fusarium* |
| GH32 | | | | OG0008649 | | OG0008649 | *<br>* | *<br>* | OG0008649 | OG0008649 | OG0008649 | OG0008649 | OG0008649<br>OG0008649<br>OG0008649 | OG0008649 | OG0008649 | OG0008649 | |
| | | | | | | OG0012516 | OG0012516 | OG0012516 | OG0012516 | OG0012516 | OG0012516 | OG0012516 | OG0012516<br>OG0012516<br>OG0012516 | OG0012516 | | | |
| | | | | OG0012851 | | * | OG0012851 | OG0012851 | OG0012851 | * | OG0012851 | OG0012851 | OG0012851 | OG0012851 | | | |
| GH78 | | | | | | * | OG0013475 | OG0013475 | OG0013475 | | * | * | OG0013475 | OG0013475 | | | |
| | | | | | | | | * | * | * | OG0014891 | OG0014891 | | | | | |
| GT1 | | | OG0013486 | | | * | OG0013486 | OG0013486 | OG0013486 | OG0013486 | OG0013486 | OG0013486 | | | | | *Fremyella diplosiphon* |
| | | | | | | | OG0014741 | OG0014741 | OG0014741 | OG0014741 | | | OG0014741 | | | | |
| | | | | | | | OG0014743 | OG0014743 | OG0014743 | OG0014743 | | | OG0014743 | | | | *Hormonema carpetanum* |
| GH3 | | | | | | | OG0015465 | OG0015465 | OG0015465 | OG0015465 | | | | | | | |
| GT2 | | | | | | | OG0014861 | OG0014861 | OG0014861 | OG0014861 | | OG0014861 | | | OG0014861 | | |
| GH43_11 | | | | | | | OG0016241 | OG0016241 | OG0016241 | | | | | | | | |
| GT21 | | | | | | | OG0016232 | OG0016232 | OG0016232 | | | | | | | | *Pyricularia oryzae* |
| GH43_24 | | OG0013753 | | | | OG0013753 | OG0013753 | OG0013753 | * | OG0013753 | | | OG0013753 | | | | *Plantactinospora* |
| CBM13 | | OG0013753 | | | | OG0013753 | OG0013753 | OG0013753 | * | OG0013753 | | | OG0013753 | | | | *Streptomyces* |
| CBM35 | | OG0013753 | | | | OG0013753 | OG0013753 | OG0013753 | * | OG0013753 | | | OG0013753 | | | | |
| GH5_16 | | | | | | * | OG0013130 | OG0013130 | * | OG0013130 | * | * | OG0013130 | OG0013130 | | | |
| GT0 | | | | | | | OG0014861 | OG0014861 | * | OG0014861 | | OG0014861 | | | | | |
| GH88 | | | | | | | | OG0018497 | | OG0018497 | | | | | | | *F. fujikuroi* |
| CE16 | | OG0015193 | | | | | OG0015193 | OG0015193 | | OG0015193 | | | | | | | |
| CBM1 | | OG0015193 | | | | | OG0015193 | OG0015193 | | OG0015193 | | | | | | | |
| GH134 | | | | | | | OG0016247 | OG0016247 | | * | | | | | | | *Pseudogymnoascus* |
| CE5 | | | | | | | OG0014836 | *<br>*<br>OG0014836 | | * | | | | | | | *F. fujikuroi* |
| CBM42 | | OG0013419 | | | | OG0013419 | | | OG0013419 | OG0013419 | OG0013419 | OG0013419 | | | OG0013419 | OG0013419 | |
| GH43_26 | | OG0013419 | | | | OG0013419 | | | OG0013419 | OG0013419 | OG0013419 | OG0013419 | | | OG0013419 | OG0013419 | |
| GH67 | | | | | OG0014678 | | | | | OG0014678 | OG0014678 | OG0014678 | OG0014678 | | | | |
| GT22 | | | | | | | * | * | * | * | OG0014367 | OG0014367 | | | | | |
| CBM67 | | | | | | | | * | * | * | OG0014891 | OG0014891 | | | | | *Streptomyces* |
| CE8 | | | | | | | | | | | OG0018533 | OG0018533 | | | | | *F. fujikuroi* |
| GH18 | | | | | | | | | | | OG0014180 | OG0014180 | OG0014180<br>OG0014180<br>OG0014180<br>OG0014180 | | | | *Trichoderma / F. fujikuroi* |
| PL1_4 | | | | | | OG0016212 | | | | | OG0016212 | OG0016212 | | | | | *F. fujikuroi* |

Supplementary table 6. The total length in base pairs and Megabase pairs with the percentage compared to the total length of these scaffold groups (table 1) of three different types of transposons (LTR, long-terminal repeats; DNA transposons; and Impalas and miniature impalas) across the different scaffold groups (core chromosomes; scaffolds specific to *F. xylarioides* and *F. udum*; *F. xylarioides*-specific scaffolds; lineage-specific scaffolds)

| | | | | | | % | | | |
|---|---|---|---|---|---|---|---|---|---|
| | | Core chromosomes | FXU | FX | LS | Core chromosomes (%) | FXU (%) | FX (%) | LS (%) |
| LTR (Mb) | Coffea674 | 1.62 | 0.74 | 0.00 | 0.74 | 3 | 17 | 3 | 40 |
| | Coffea659 | 1.03 | 1.19 | - | 0.17 | 2 | 15 | | 8 |
| | Robusta277 | 1.45 | 1.38 | 0.02 | 0.27 | 3 | 18 | 34 | 12 |
| | Robusta254 | 1.18 | 1.15 | 0.01 | 0.21 | 2 | 16 | 33 | 12 |
| | Arabica563 | 1.52 | 1.19 | 0.00 | 0.18 | 3 | 13 | 9 | 9 |
| | Arabica908 | 1.90 | 1.21 | 0.03 | 0.28 | 4 | 16 | 5 | 13 |
| DNA (Mb) | Coffea674 | 1.60 | 0.43 | 0.00 | 0.18 | 3 | 10 | 3 | 10 |
| | Coffea659 | 1.09 | 0.40 | - | 0.27 | 2 | 5 | | 12 |
| | Robusta277 | 0.98 | 0.23 | 0.01 | 0.21 | 2 | 3 | 13 | 10 |
| | Robusta254 | 0.85 | 0.19 | 0.00 | 0.16 | 2 | 3 | 0 | 9 |
| | Arabica563 | 1.03 | 0.25 | 0.01 | 0.23 | 2 | 3 | 17 | 11 |
| | Arabica908 | 0.95 | 0.34 | 0.01 | 0.10 | 2 | 4 | 1 | 5 |
| Impala and mimps (bp) | Coffea674 | 45,269 | 14,248 | | 28,032 | 0.1 | 0.3 | - | 1.5 |
| | Coffea659 | 45,612 | 14,248 | | 67,808 | 0.1 | 0.2 | | 3.1 |
| | Robusta277 | 37,858 | 7,027 | | 15,181 | 0.1 | 0.1 | - | 0.7 |
| | Robusta254 | 48,706 | 1,569 | | 886 | 0.1 | 0.0 | - | 0.0 |
| | Arabica563 | 38,534 | 15,231 | | 42,830 | 0.1 | 0.2 | | 2.0 |
| | Arabica908 | 38,814 | 7,531 | | 7,982 | 0.1 | 0.1 | | 0.4 |
| All transposons (LTRs, DNAs, Impalas) | Coffea674 | 3.3 | | | | 6 | 27 | | 51 |
| | Coffea659 | 2.2 | | | | 4 | 15 | | 23 |
| | Robusta277 | 2.5 | | | | 5 | 16 | | 22 |
| | Robusta254 | 2.1 | | | | 4 | 17 | | 21 |
| | Arabica563 | 2.6 | | | | 5 | 13 | | 21 |

Supplementary table 7. Accession numbers and source details for each Impala and miniature Impala (mimps) (continued overleaf)

| Accession | Transposon | Sequence |
|---|---|---|
| AF076624.1 | F. o. repetitive element mimp1 | |
| AF076625.1 | F. o. repetitive element mimp2 | |
| EU833100.1 | F. o. f. sp. melonis MITE mimp3 | complete sequence |
| EU833101.1 | F. o. f. sp. lycopersici MITE mimp4 | complete sequence |
| AF282722.1 | F. o. f. sp. melonis transposon impala transposase gene | complete cds |
| AF363394.1 | F. o. f. sp. bulbigenum translation factor EF1 alpha gene | partial cds |
| AF363395.1 | F. o. f. sp. melonis translation factor EF1 alpha gene | partial cds |
| AF363396.1 | F. o. f. sp. vasinfectum translation factor EF1 alpha gene | partial cds |
| AF363397.1 | F. o. f. sp. melonis translation factor EF1 alpha gene | partial cds |
| AF363398.1 | F. o. f. sp. melonis translation factor EF1 alpha gene | partial cds |
| AF363399.1 | F. o. f. sp. lycopersici translation factor EF1 alpha gene | partial cds |
| AF363400.1 | F. o. f. sp. melonis translation factor EF1 alpha gene | partial cds |
| AF363401.1 | F. o. f. sp. radicis-lycopersici translation factor EF1 alpha gene | partial cds |
| AF363402.1 | F. o. f. sp. ciceri translation factor EF1 alpha gene | partial cds |
| AF363403.1 | F. o. f. sp. melonis translation factor EF1 alpha gene | partial cds |
| AF363404.1 | F. o. f. sp. lini translation factor EF1 alpha gene | partial cds |
| AF363405.1 | F. o. f. sp. lini translation factor EF1 alpha gene | partial cds |
| AF363406.1 | F. o. f. sp. dianthi translation factor EF1 alpha gene | partial cds |
| AF363407.1 | F. o. f. sp. melonis transposon impala M24-impE | partial sequence |
| AF363412.1 | F. o. f. sp. lini transposon impala Ln3-1 | partial sequence |
| AF363413.1 | F. o. f. sp. lini transposon impala Ln88-23 | partial sequence |
| AF363414.1 | F. o. f. sp. cubense transposon impala Cu-12 | partial sequence |
| AF363416.1 | F. o. f. sp. phaseoli transposon impala Ph-5 | partial sequence |
| AF363417.1 | F. o. f. sp. phaseoli transposon impala Ph-9 | partial sequence |
| AF363418.1 | F. o. f. sp. albedinis transposon impala A-33 | partial sequence |
| AF363419.1 | F. o. f. sp. soil transposon impala S47-35 | partial sequence |
| AF363420.1 | F. o. f. sp. raphani transposon impala R-8 | partial sequence |
| AF363425.1 | F. o. f. sp. melonis transposon impala M24-impD | partial sequence |
| AF363426.1 | F. o. f. sp. melonis transposon impala MK14 | partial sequence |
| AF363427.1 | F. o. f. sp. lini transposon impala Ln88-8 | partial sequence |
| AF363428.1 | F. o. f. sp. radicis-lycopersici transposon impala RL28delta22 | partial sequence |
| AF363429.1 | F. o. f. sp. melonis transposon impala MKdelta208 | partial sequence |
| AF363430.1 | F. o. f. sp. lycopersici transposon impala L15delta5 | partial sequence |
| AF363432.1 | F. o. f. sp. lycopersici transposon impala L15-15 | partial sequence |
| AF363433.1 | F. o. f. sp. radicis-lycopersici transposon impala RL28-17 | partial sequence |
| AF363434.1 | F. o. f. sp. lini transposon impala Ln86-10 | partial sequence |
| AF363435.1 | F. o. f. sp. ciceris transposon impala Ci-36 | partial sequence |
| AF363436.1 | F. o. f. sp. ciceris transposon impala Ci-16 | partial sequence |
| AF363437.1 | F. o. f. sp. melonis transposon impala MK28 | partial sequence |

Supplementary table 6 continued

| | | |
|---|---|---|
| AF363438.1 | F. o. f. sp. lycopersici transposon impala L15-16 | partial sequence |
| AJ608703.3 | F. o. f. sp. lycopersici shh1 gene | |
| AJ608703.3 | F. o. f. sp. lycopersici fot5 gene | |
| JX204302.1 | F. o. f. sp. fragariae transposon Impala_1 | complete sequence |
| Schmidt et al. 2013 | FoCrypton | |
| Schmidt et al. 2013 | FoHelitron | |
| Schmidt et al. 2013 | Fot6 | |
| Schmidt et al. 2013 | Fot8 | |
| Schmidt et al. 2013 | Hop3 | |
| Schmidt et al. 2013 | Hop6 | |
| Schmidt et al. 2013 | MGR583-like | |
| Schmidt et al. 2013 | Nht2-like | |
| Schmidt et al. 2013 | YahAT4 | |
| Schmidt et al. 2013 | YahAT6 | |
| Schmidt et al. 2013 | Yaret1 | |
| Schmidt et al. 2013 | Yaret2 | |
